# Incorporating temporal information during feature engineering bolsters emulation of spatio-temporal emergence

**DOI:** 10.1101/2024.01.11.575245

**Authors:** Jason Y. Cain, Jacob I. Evarts, Jessica S. Yu, Neda Bagheri

## Abstract

**Motivation:** Emergent biological dynamics derive from the evolution of lower-level spatial and temporal processes. A long-standing challenge for scientists and engineers is identifying simple low-level rules that give rise to complex higher-level dynamics. High-resolution biological data acquisition enables this identification and has evolved at a rapid pace for both experimental and computational approaches. Simultaneously harnessing the resolution and managing the expense of emerging technologies—e.g. live cell imaging, scRNAseq, agent-based models—requires a deeper understanding of how spatial and temporal axes impact biological systems. Effective emulation is a promising solution to manage the expense of increasingly complex high-resolution computational models. In this research, we focus on the emulation of a tumor microenvironment agent-based model to examine the relationship between spatial and temporal environment features, and emergent tumor properties.

**Results:** Despite significant feature engineering, we find limited predictive capacity of tumor properties from initial system representations. However, incorporating temporal information derived from intermediate simulation states dramatically improves the predictive performance of machine learning models. We train a deep-learning emulator on intermediate simulation states and observe promising enhancements over emulators trained solely on initial conditions. Our results underscore the importance of incorporating temporal information in the evaluation of spatio-temporal emergent behavior. Nevertheless, the emulators exhibit inconsistent performance, suggesting that the underlying model characterizes unique cell populations dynamics that are not easily replaced.

**Availability:** All source codes for the agent-based model, emulation, and analyses are publicly available at github.com/bagherilab/ARCADE, github.com/bagherilab/emulation, and github.com/bagherilab/emulation_analysis, respectively.

## Introduction

Grand challenges in biology have required tools with increasingly higher resolution and throughput while also sampling across spatial and temporal axes (Lähnemann et al., 2020; Eftimie, 2022; Bindea et al., 2013; Johnson et al., 2021). In particular, dynamic data from live cell imaging is positioned to become the next “omics” (Lelek et al., 2021; Bagheri et al., 2022) with lower-level resolution data, like scRNAseq, contributing a more nuanced understanding of individual system components across space (Hwang et al., 2018; Heumos et al., 2023). Our ability to utilize high resolution data has often lagged behind our ability to generate said data (Ji et al., 2017; Lähnemann et al., 2020; Eftimie, 2022). Parity between computational and experimental approaches will allow researchers to synergistically utilize computational models to explain nonintuitive observations, identify “rules of life”, and design model-driven experiments that test new hypotheses. This parity will identify both temporal and spatial components that are fundamental to specific biological systems *a priori* (Müller and Pörtner, 2021; Eftimie et al., 2023). In order to address this knowledge gap, we interrogate the use of statistical emulation to estimate emergent behavior in a comprehensive analysis of a high-resolution agent-based model.

Computational models capable of characterizing cellular dynamics over time and space across multiple scales are fundamental to scientific progress. Agent-based models (ABMs) have gained popularity due to their ability to simulate populations of heterogenous agents dynamically over time and in space in order to predict system-level emergent properties (Shi et al., 2014; Vodovotz and An, 2019; West et al., 2023). ABMs are designed using the behavior and interactions of individual agents (usually cells) in an evolving spatial and temporal context (Bonabeau, 2002; Sklar, 2007). This rule-based approach is well suited to modeling emergent behaviors, such as the development of multi-cellular systems (Ghaffarizadeh et al., 2018; Osborne et al., 2017; Corti et al., 2021; Pleyer and Fleck, 2023) or the spread of infectious diseases (Hunter et al., 2018; Kerr et al., 2021; Gomez et al., 2021). ABMs can capture the heterogeneity and the stochasticity of biological systems, as well as the bilateral relationship between the local microenvironment and agents’ behaviors (Metzcar et al., 2019; Yu and Bagheri, 2021; Norton et al., 2019). Furthermore, ABMs can be used to investigate the effects of different parameters and conditions on emergent behaviors, providing insights into complex biological processes that may be difficult or impossible to observe experimentally (Yu and Bagheri, 2016; Virgilio et al., 2018; Yu and Bagheri, 2021; Soheilypour and Mofrad, 2018; Peng and Vermolen, 2020). As such, ABMs are an increasingly important—albeit computationally expensive—tool for understanding complex biological systems and generating testable hypotheses that can inform experimental design.

Statistical emulation (SE), an approach to surrogate modeling, generates statistical models by mapping independent variables (inputs) to dependent variables (outputs) via statistical inference and machine learning (ML) with limited knowledge of the underlying simulation model. Effective SE produces simplified models that are computationally cheaper to analyze, and therefore interrogate, relative to the original model. The flexible pattern recognition of ML provides two key benefits: (1) the emulation model building process codifies selection of requisite inputs to identify dominant statistical patterns; and (2) the mathematical frameworks involved in pattern recognition—once trained—are easily calculated via sequences of simple mathematical operations. On the contrary, the feature selection and design process required for SE is laborious, and the algorithms demand significant data for adequate performance. Balancing these trade-offs supports effective integration into multi-class models (Cicchese et al., 2017; Cess and Finley, 2020), sensitivity analyses for complex models (Alden et al., 2020), and parameter sweeps of mechanistic models (Vernon et al., 2018; Wang et al., 2019). Even with these successes, there have been few comprehensive analyses of the application of emulation models to rule-based models designed to simulate tissue dynamics that emerge from temporal and spatial interactions (Alden et al., 2020; Kieu et al., 2022; Angione et al., 2022). SE of ABMs is largely under-explored despite knowing that the computational demands of large multi-scale ABMs can outpace their utility to generate hypotheses between continuous input variables and heterogeneous outputs (Glen et al., 2019; Heppenstall et al., 2021). Synergistic development of SE frameworks for ABMs would facilitate expedited analyses and provide systematic methods towards hypothesis generation. Understanding the characteristics (e.g. spatial, temporal, etc.) of data required to emulate key ABM dynamics provides a more thorough understanding of the drivers of emergent outcomes.

### The tumor microenvironment as a model system

ABMs of the tumor microenvironment have leveraged *in silico* networks to represent the vascular environment (Norton et al., 2018; Metzcar et al., 2019; Yu and Bagheri, 2021). Physical and structural characteristics (e.g. pressure, shear, radius) are often encoded into network elements like edges and nodes (Fredrich et al., 2018, 2019). Networks are flexible data structures that can be modified and interrogated, enabling researchers to study interplay among agents and between agents and environments. Functional coupling between the agents and the environment is required to capture experimentally observable divergent emergence like vascular collapse and necrotic core formation (Norton et al., 2018; Yu and Bagheri, 2021).

Network analysis provides a means to abstract high-resolution, spatial information into summary statistics (Pavlopoulos et al., 2011; Koutrouli et al., 2020). Network topology and morphology have been used to understand ecological systems (Peterson et al., 2013; Modica et al., 2021), interrogate biological pathways (Magnano and Gitter, 2021), analyze neurological structure (Sporns, 2013; Kok et al., 2020; Bassett and Sporns, 2017), and identify novel treatment (Iadevaia et al., 2010). Specifically in tumor development, network analyses are a promising approach to study healthy and pathogenic vascular mimicry and angiogenesis (Amat-Roldan et al., 2015; Alves et al., 2018; Fouladzadeh et al., 2021). Thus, we hypothesize that vascular network analysis enables the encoding of complex network architecture as interpretable inputs for SE.

In this study, we utilize network analysis to construct SE models of ARCADE, an existing agent-based model of a tumor microenvironment with heterogeneous and realistic vascular networks (Yu and Bagheri, 2021). Our investigation reveals a limited relationship between spatial network topology characteristics and emergent tumor properties. We demonstrate the efficacy of incorporating temporal evolution of network metrics to predict emergent properties of the system. Leveraging this temporal information, we develop deep-learning models that improve the predictive power of emulators. Our results highlight the role of temporal dynamics in understanding and predicting emergent properties of diverse cell populations that evolve from lower-level spatial and temporal processes.

## Materials and methods

### Data and code availability

All source code for the ARCADE ABM is publicly available on GitHub at github.com/bagherilab/ARCADE. The scripts used to perform analyses and generate figures reported in this paper are also publicly available on GitHub at github.com/bagherilab/emulation_permutation.

### ARCADE

All ABM simulations and model analyses were performed similar to those described in a previous publication (Yu and Bagheri, 2021) using ARCADE v2.4 (Yu, 2023). ARCADE is an on-lattice, agent-based model designed to represent tissue and tumor development. Agents in ARCADE represent tissue cells that can take on seven cell states—quiescent, migratory, proliferative, apoptotic, necrotic, senescent, and undecided. A cell’s transition between states is determined by accounting for their current properties (e.g. age, size), relationship to other agents (e.g. crowding), and local environmental conditions (glucose, oxygen, and TGF*α*). Each agent contains a metabolism module tracking the conversion of nutrients into energy and cell mass, and a signaling module comprising a ligand-sensing network guiding cell decisions between proliferative and migratory states. For instance, the intracellular signaling state impacts the individual cell’s probability between proliferation and migration. The resulting cellular states determine the discrete behavior performed by the cells at each simulation tick (one minute) within a hexagonal grid. A more detailed description of agent states, rules, and modules can be found in Yu and Bagheri (2020).

The simulation environment is described by a higher resolution triangular lattice containing the concentration of molecules (oxygen, glucose, and TGF*α*). The molecules diffuse through the environment according a diffusion partial differential equation model given by:

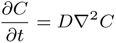

The dynamic vasculature model is represented as a graph data structure, utilizing graph embeddings to represent hemodynamic properties (shear stress, circumferential stress, and flow rate) of the capillaries. These properties are derived from modeling capillary flow through each individual edge with fixed conditions at the originating artery and vein roots. Vascular structure and function change over time based on their coupling with cell agent populations. Functional stresses from tissue cells lead to vascular remodeling, changing vessel radius and wall thickness as a function of hemodynamic properties and metabolic demand. Cancer cells degrade the vasculature by removing components; this removal can lead to larger scale disruption of vascular structure, as functional vasculature requires perfusion. A more detailed description of the environment, dynamic vasculature initialization, and subsequent physical modeling can be found in Yu and Bagheri (2021).

Two agent populations, healthy and cancer cells, exist in the simulations. *In silico* cancer cell populations exhibit hallmarks of cancer: increased crowding tolerance, increased preference for metabolic glycolysis than oxidative phosphorylation, and increased migratory invasiveness versus healthy cell populations (Hanahan and Weinberg, 2011; Vander Heiden et al., 2009). Towards feasibility and reproducibility, this study does not account for healthy cells evolving to take on cancer hallmarks. The colony and tissue contexts describe simulations comprised of solely cancer cells and a combination of cancer and healthy cell populations, respectively. Prior work highlighted differences in emergent behavior between colony and tissue contexts (Yu and Bagheri, 2016, 2021; Prybutok et al., 2022b).

### ARCADE simulations and workflow

We focus on emulating three emergent tumor properties: activity (instantaneous state of system), growth rate (cumulative temporal behavior), and symmetry (instantaneous spatial state). Activity is the ratio of active to dead cells, describing the degree of necrosis in the tumor. Growth rate is a temporal emergent property that generally describes tumor aggression. Symmetry is a spatial emergent property describing the density and implying the aggressiveness of the tumor. Specific calculations for these properties are detailed in Supplemental information.

The initial vascular structure is the only differing variable between simulations of the same context. Vasculatures are stochastically generated using starting root geometries, detailed in a previous article (Yu and Bagheri, 2021). 100 seeds generated a unique vasculature for each starting root geometry and seed combination.

### Network analysis and feature extraction

To represent the intricate characteristics of vasculatures within ABM simulations, we employed network analysis. By using a graphical representation of the vasculature, we utilized graph theory metrics as structural and functional features for our ML models. Graph theory provides a number of benefits: it comprises diverse metrics that account for the topological structure of the vasculature; it has mechanisms for specifying vessel importance; and it can quantify overall vessel connectivity. Vascular structure is represented as a network where blood vessels are represented as edges, and where junctions and end points are represented as nodes.

We quantify properties of the *in silico* vasculature and use these features to predict spatial and temporal emergent dynamics (Figure 4). Aggregate hemodynamics, such as flow and wall thickness, are calculated for each vessel segment and then averaged. Topological features are calculated using the igraph package (Csardi and Nepusz, 2005) in Python to create a network representation of the vasculature. Hemodynamic edge features are the same network metrics used to define topological features with one modification: edges in the graph are weighted by hemodynamic properties. Finally, spatial features use distance from the center of the simulation (the tumor seeding location) as edge weights, such that vessels within and closer to the tumor core are weighted more heavily. The supplementary materials (Supp. Table 1, 2, and 3) offer detailed information about specific graph metrics and how they were obtained.

### Statistical emulation

#### Machine learning models and hyperparameter selection

Python package sklearn was used to build the ML models (Pedregosa et al., 2011). All hyperparameters specified are referred to as their respective arguments for each model.

A Sobol random search (Sobol, 1976) was used to select the tested hyperparameters during cross-validation using the scipy (Virtanen et al., 2020) package in Python. The ranges used for the Sobol random search are detailed in Table 1. Parameter types categorized as “linear” used Sobol indices in the linear space, whereas those types categorized as “logarithmic” used the log of the bounds as the search range. Every discrete parameter value was exhaustively tested in combination with parameter values selected using a Sobol random search on continuous parameter spaces. Each set of Sobol and discrete hyperparameters was used to generate an independent ML model. The model with the best average performance metric (we use the coefficient of determination *R*^2^) compared across all hyperparameters during cross-validation was used for training and testing. 10-fold cross-validation was used in all cases. The coefficient of determination is defined as 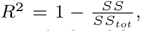, where *SS* is the sum of squares of residuals ML model and *SS_tot_* is the total sum of squares from the mean. *R*^2^*_val_* was calculated from ML models evaluated on the validation data set. The reported *R_train_*^2^ and *R*^2^*_test_* were calculated on the training and withheld testing sets after cross-validation. The necessary training times to complete hyperparameter selection are included in Supp. Table 4.

**Table 1.** Continuous hyperparameters used in cross-validation Sobol search.

|  | Hyperparameter | Bounds |  | Type |
| --- | --- | --- | --- | --- |
|  |  | Lower | Upper |  |
| MLR | alpha | 0.001 | 1.000 | Log |
|  | l1_ratio | 0.1 | 1.0 | Linear |
| SVR | C | 0.0001 | 1.000 | Log |
|  | epsilon | 0.0 | 1.0 | Linear |
| RF | n_estimators | 1 | 100 | Linear |
|  | max_features | 0.01 | 1.0 | Log |
|  | max_depth | 1 | 100 | Linear |
|  | min_samples_split | 0.01 | 1.0 | Log |
|  | max_samples_leaf | 0.01 | 1.0 | Log |
| MLP | alpha | 0.0001 | 1.000 | Log |

**Table 2.**
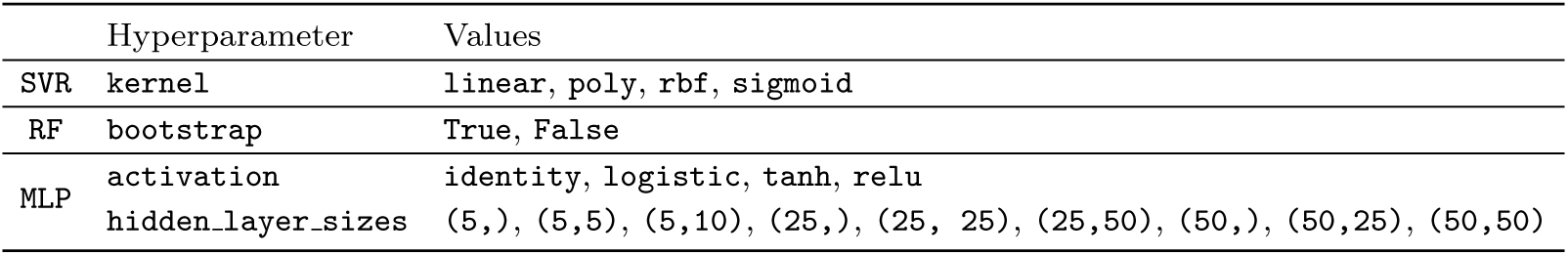
Discrete hyperparameters used in cross-validation.

#### Recurrent neural network architecture and training

In order to account for dynamic network evolution, we trained deep neural networks (NNs) to use time course information to improve predictive performance using the keras (Chollet et al., 2015) API of the tensorflow (Abadi et al., 2015) package in Python. The purpose of the trained NNs was to predict network evolution from the initial vascular structure in order to increase prediction performance of the emulation models. The neural network utilized a long short-term memory (LSTM) layer, a type of recurrent NN layer capable of capturing sequential patterns (Hochreiter and Schmidhuber, 1997). The LSTM layer was followed by 3-4 fully connected layers. Full network topology details are described in Supp. Figure 10.

Network topology features from each simulation day were collected and stacked into multivariate time series to facilitate transfer learning of NN parameters. The recurrent NN (RNN) was pre-trained on the full-length time series, encompassing the entire temporal evolution of the network in order to constrain the RNN parameters. To further fine tune the RNN, bootstrapped samples from a subset of the time series (10 days, 5 days, 3 days) were used to sequentially retrain the model (using the previous pretrained deep-learning models as a starting point) and retrain on the initial conditions to provide the ultimate emulator. We applied the best performing model to predict the two week network structures for a reserved test set from only the initial network topology, and then passed the predicted features into the naive ML models (Figure 4A).

## Results

The objective was to predict the emergent tumor output metrics by their respective simulation inputs using naive ML algorithms (Figure 1). Specifically, we used multiple linear regression (MLR), random forest (RF), support vector regression (SVR), and multi-layer perceptron (MLP) to predict emergent tumor output metrics (activity, growth, and symmetry) from the initial condition of the microenvironment vasculature. In general, we use “emulators” to describe those models accepting initial conditions as the sole inputs and we us “ML models” to describe those accepting any other timepoint. The microenvironment was represented by a network analysis of the vascular architecture to provide features for ML models.

**Fig. 1:**
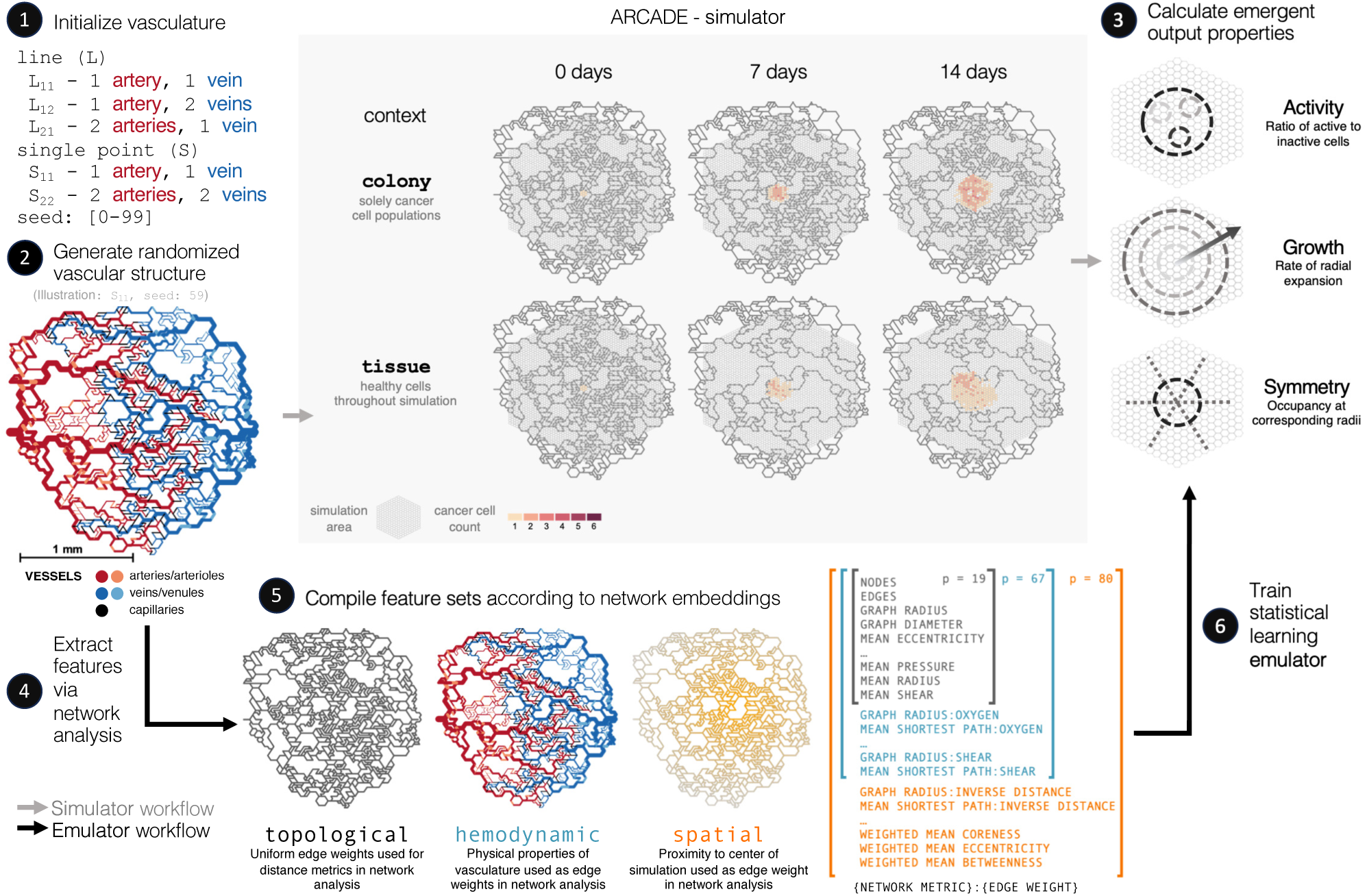
Emulation workflow. A summary of the overall emulation workflow. 1) Vasculature structures are generated based on the a starting root geometry (single point vs. line roots, and the number of initializing arteries and veins) and a random seed, where 0-99 were used. 2) ARCADE, an ABM of the tumor microenvironment, receives *in silico* vasculature networks and initial cell population colonies as inputs. ARCADE simulates intra- and inter-cellular interactions among diverse agents to predict the evolution of vascular architecture and function, as well as the evolution of cell populations, over space and time. Two different simulation contexts were used to initialize populations: colony and tissue. 3) Spatio-temporal dynamics are summarized with output metrics that evaluate emergent tumor properties at the end of the simulations: activity, growth, and symmetry. 4) Network metric-based feature sets are extracted from vascular architectures. Nodes represent junctions in the vasculature; edges represent sources of nutrients in the simulation. 5) Feature sets are aggregated based on the information used. Topological features are extracted from the unweighted structure of the network. Hemodynamic features are extracted from attributes of network topologies including hemodynamic characteristics as edge weights. Spatial features account for distance between the information in the network from the center of the simulation. 6) Statistical learning models use network metric-based feature sets to predict emergent tumor output metrics.

Various network analyses generated a suite of feature sets that were used to train the ML models. The topological feature set characterized traditional structural and topological information of the vasculature system (e.g. eccentricity, betweenness, average degrees), as well as mean hemodynamics across the vascular system (e.g. mean pressure, mean shear, mean radius, etc.). The hemodynamic feature set ascribed hemodynamics properties (e.g. flow, pressure, wall thickness) as edge weights to the topological features in order to capture vascular function. The spatial feature set integrated relative locations of edges and nodes from the center of the simulation—which represents the center of the tumor—and accounted for these properties as both additional edge weights and penalties in weighted average calculations. Each feature set is inclusive of the previous set: topological *⊆* hemodynamic *⊆* spatial. A comprehensive feature set breakdown is described in Supp. Tables 1, 2, and 3.

### Unweighted and hemodynamic-weighted network topologies do not predict emergent tumor output metrics

In order to represent vascular structures in a quantitative format, topological network data from initialization vasculatures were used to train the emulators. Aggregate network metrics (e.g. number of nodes and edges) characterized the size and density of the network. Additional metrics—e.g. average eccentricity, betweenness, and coreness of each node—were used to characterized the average behavior of nodes in the network. This topological feature set does not include spatial node embeddings as a factor in the analysis.

Vascular structures represented by initial topological features are not predictive of emergent behavior, resulting in models that exhibit both overfit and underfit characteristics. The coefficient of determination for test data in all topological emulation models is less than 0.3 (Figure 2A), suggesting that these models are underfit as a result of the variance in our features not explaining the variance in the emergence. The substantial performance gaps between training and testing results that derive from more complex algorithms (i.e. SVR, MLP) indicate overfitting, despite implementing regularization (Figure 2A). The best cross-validation performance corresponded with better training performance in simpler algorithms (i.e. MLR, RF), whereas the large majority of cross-validation models had better training metrics than the selected model in the more complex algorithms (Supp. Figure 1). We find that the contribution of each feature— based on a permutation importance analysis—was inconsistent between emulators, likely resulting from the collinearity of features and the degree of regularization required for more complex ML algorithms 2).

**Fig. 2:**
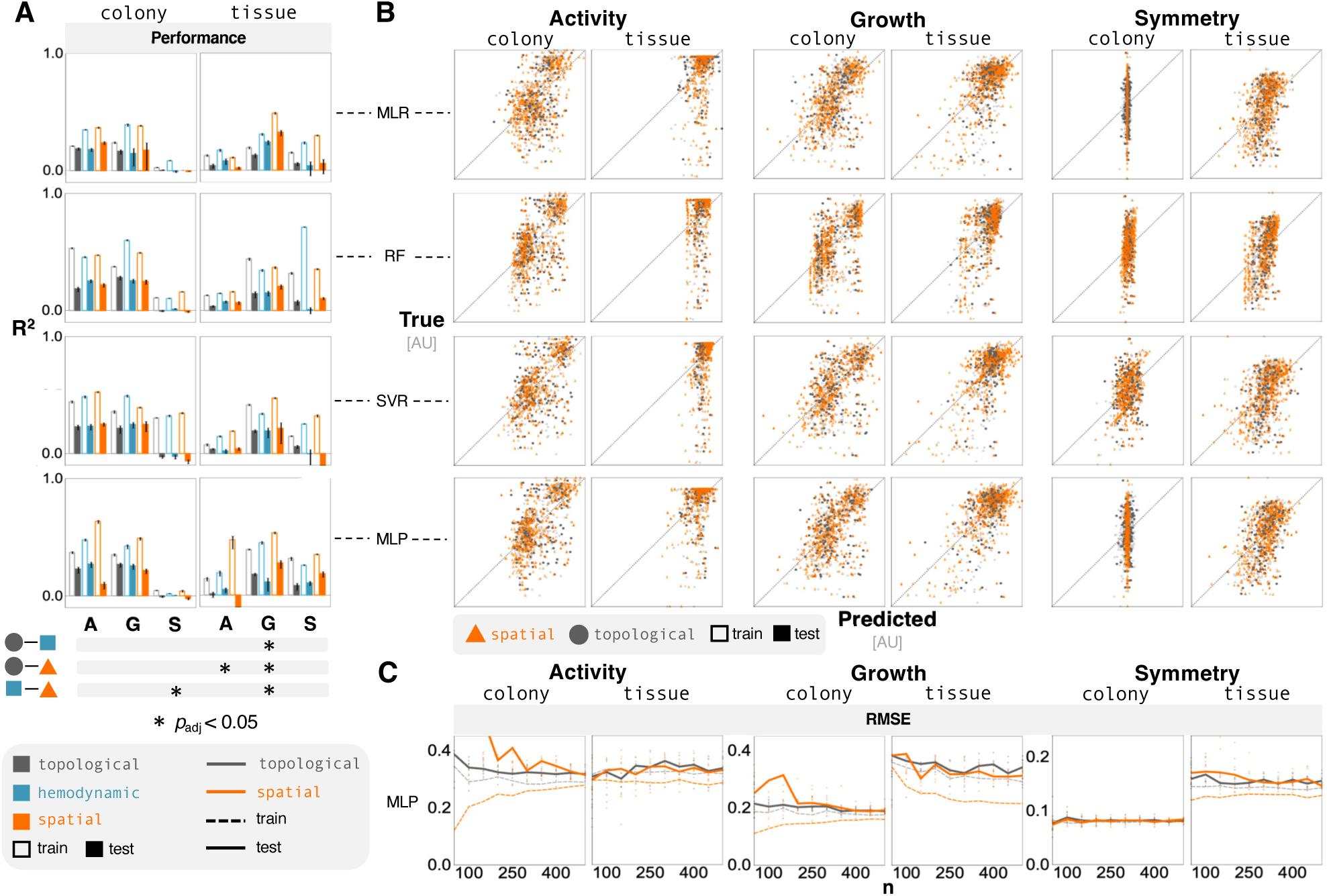
Spatial information does not support emulation. (**A**) Bar plots show predictive performance of emergent outputs (A: activity, G: growth, S: symmetry) across feature sets for different models (MLR, RF, SVR, and MLP). Feature engineering offered limited improvement. Bar chart values range from -0.1 to 1.0; the horizontal axis is at 0.0. The Bonferroni corrected p-values from a two-way ANOVA highlight significant results (noted with asterisks) that have an adjusted p-value less that 0.05. (**B**) Parity plots show differences between the variance in the predicted response and the true response, comparing the topological and spatial feature sets. (**C**) Additional training data offered diminishing returns on predictive performance of MLP models that were trained on both spatial and topological features. These subplots show the average RMSE as a function of the size of training data. The individual points represent the RMSE from randomized test sets.

Performance is variable as a result of context; activity predictions in colony contexts (Figure 2A, left) performed more accurately than comparable models in tissue contexts (Figure 2A, right). On the contrary, emulators that comprise healthy cell background in the tissue context (Figure 2A, right) reflect more accurate symmetry and growth predictions. Notably, an adequate model—exhibiting a test coefficient of determination over 0.0—for symmetry in the colony context could not be trained.

We then included hemodynamic characteristics as edge weights in the network analysis for network-distance metrics in the distance-based analyses. These hemodynamic features describe higher-resolution physical characteristics of the environment that have clinical implications. For example, pressure-based metrics have been associated with system-level hypertension. (Blanco et al., 2017) The resulting weighted network analysis metrics offered limited improvement in prediction accuracy; most results were statistically insignificant relative to the unweighted case, determined by a two-way ANOVA (Figure 2A). Training and testing performance of hemodynamic emulators experienced diminishing returns as the amount of training data increased (Supp. Figure 3).

### Network topologies with hemodynamic and spatial information do not predict emergent tumor output metrics

We hypothesized that the spatial variance in vascular structure seeds would account for a significant amount of the variance found in emergent tumor behaviors. This hypothesis was motivated by the finding that vascular collapse, a large-scale dynamic event, substantially impacts the spatial structure of simulations. The consequence of collapse in concert with vessel degradation is required to observe the formation of a necrotic tumor core (Yu and Bagheri, 2021). Thus, in order to capture spatial information in the network analysis, we applied a proportional penalty for edges and nodes based on their respective euclidean distance from the center of the simulation (a proxy for the tumor core).

We investigated whether the distance of network nodes from the center of the simulation would explain tumor behavior (Figures 2A and 2B). Including spatial features led to negligible, if any, improvement on the emulators’ testing performance. Regularization in the cross-validated emulation models led to predicted targets that spanned less variance in the response variable (Figures 2), which was indicated by the low coefficients of determination (Figure 2A). The RMSE of the predicted targets showed minimal improvement as we increased the number of training data (Figure 2C), suggesting that the poor performance was not a result of insufficient training data; instead, it likely derived from an incomplete representation of the drivers of emergence.

### Network analysis of environmental snapshots at later timepoints predict emergent tumor output metrics

In order to interrogate the impact of network evolution on the performance of our ML models, we calculated hemodynamic network metrics for the vascular structure on each simulation day. Features at each timepoint were solely derived from the snapshot at the later vascular architecture. These temporally dynamic features were then used to predict emergent tumor dynamics.

We found that network metrics that were generated from later timepoints provided better predictive performance (Figure 3A). In the colony context activity was significantly more predictable when using features representing network properties at later timepoints (Figure 3B, left). Additionally, we observed non-monotonic improvements in growth prediction with a notable improvement in performance using snapshots of network properties from the middle of the time course (Figure 3B, middle, Supp. Figure 7). Symmetry showed minimal improvement (Figure 3B, right). Conversely, in the tissue context, predictions of activity did not improve, while predictions of growth improved monotonically (Figure 3B, Supp. Figure 8). The improvement in predicting symmetry from initial features relative to features from two simulation weeks was larger in tissue contexts than in colony contexts (Figure 3B, Supp. Figure 8).

**Fig. 3:**
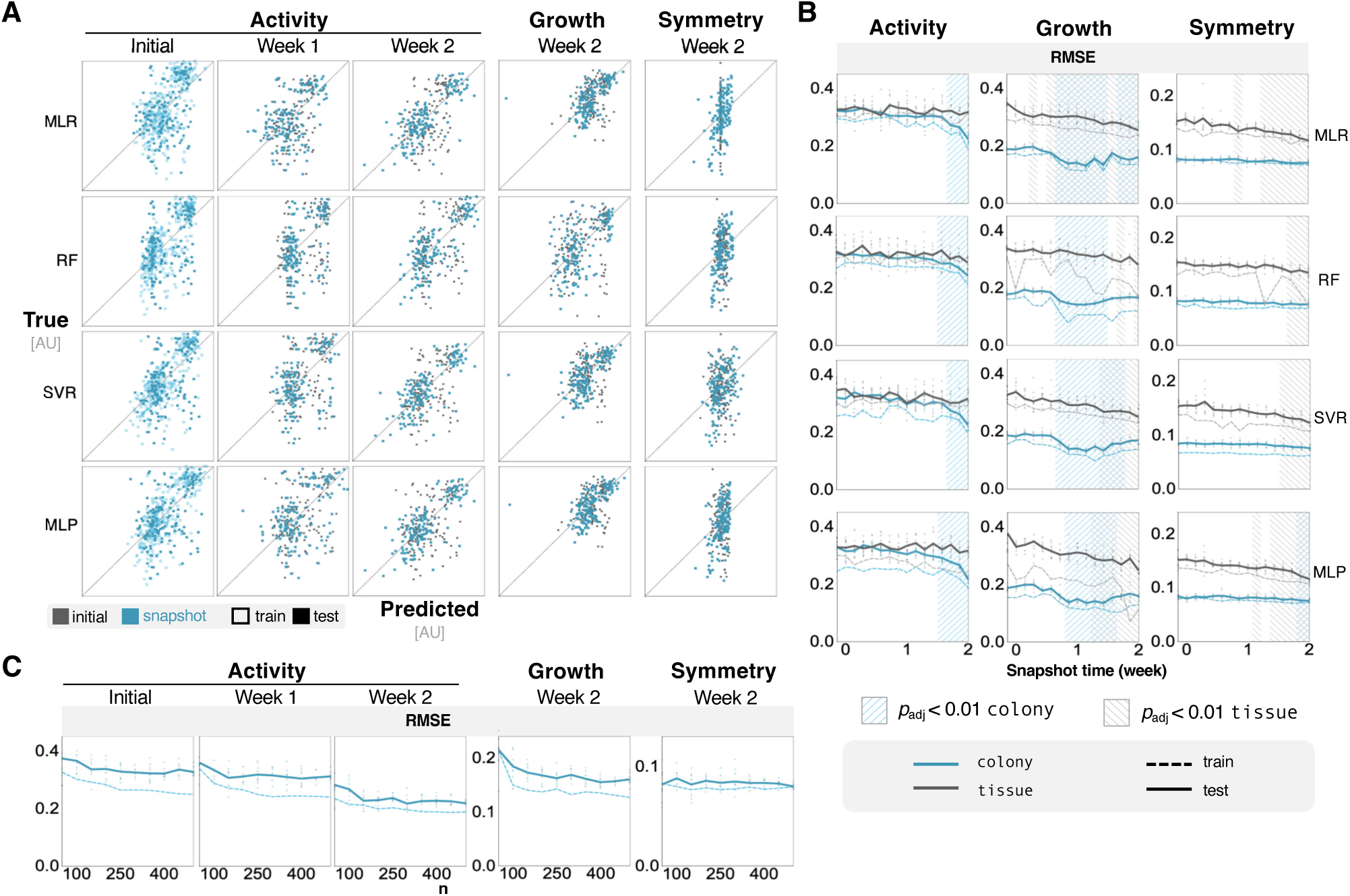
Temporal information improves accuracy of ML models. (**A**) Parity plots show the predictive performance of colony context ML models that were trained on exclusively on features from later timepoints. The trends are consistent for all three predicted outputs. The week 1 parity plots for growth and symmetry, and the corresponding results for tissue context, are included in Supp. Fig. 7, and 8. (**B**) Line plots show improvement of ML models in both colony and tissue contexts when they are trained on features from timepoints later in the simulation. One-way Dunnet (Dunnett, 1955) statistical test with Bonferroni correction show timepoints features that perform better than the initial timepoint. Hash marks signify adjusted p-values less than 0.01. (**C**) Predictive performance as a function of training data for MLP models at later timepoints. These subplots show the average RMSE as a function of the size of training data. The individual points represent the RMSE from randomized test sets.

The analysis of vascular structure included features post-vascular collapse, which likely accounted for some of the better performance. Alternatively, including differential timepoint analyses did not provide additional benefits to predictive power (Supp. Figure 4, Supp. Figure 5). Simply shortening the simulation time, and therefore the prediction horizon, to one week resulted in a sharp decline in emulator performance (Supp. Figure 6). Once again, there was limited improvement in test performance with additional training data, indicating there was sufficient data to train these ML models (Figure 3C).

### Neural network structures trained on network dynamics support ABM emulators

We hypothesized that including temporal dynamics in the training of an emulator could improve performance based on the enhanced predictive potential of later timepoints. First we trained a recurrent NN (RNN) on the network evolution of the vascular architecture in order to forecast the network metrics of the final timepoint based on the initial condition (Figure 4A). Then, we used statistical learning models to predict emergent outputs using the predicted final network metrics as features (Figure 4B).

**Fig. 4:**
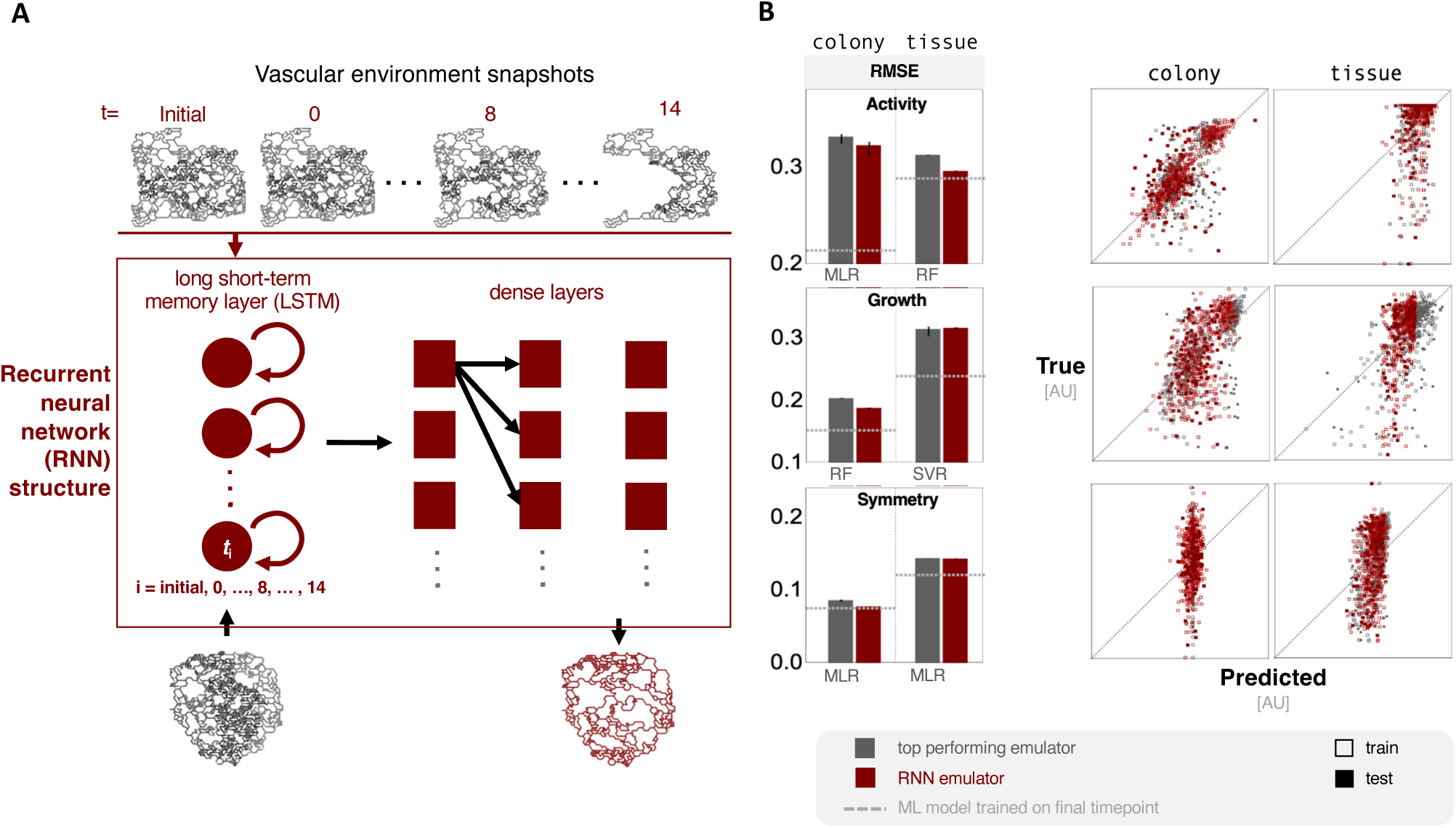
Incorporating temporal information can improve emulation performance. (**A**) A summary of the vascular network structure and ML model workflow. Network metrics from vascular structures from consecutive simulation days are used as a sequence to train the RNN. Initial vasculatures (represented by the greyscale network) are then used to predict network properties of the vasculatures after two simulation weeks (red network); the two-week predictions are then used to predict emergent behaviors with ML models. (**B**) Bar plots show performance between emulators trained on forecasted network metrics and the top performing naive emulation models (noted beneath bars). Associated parity plots show the prediction performance of the RNN models across contexts for each emergent output.

The efficacy of using forecasted features varied widely by emergent outcome and context. Emulators trained on forecasted features never performed worse than those trained on initial-condition-derived features (Figure 4B, left). In the colony context the RNN model led to a small improvement in activity and growth, while symmetry remained largely unpredictable. In the tissue context, the model was improved across all emergent behaviors. Activity predictions improved in the tissue context with a test *R*^2^ nearly three times greater than the naive emulation models (Figure 4B).

While growth exhibited the largest benefits from training on temporal dynamics, all predictions across all emergent outcomes presented at least minor improvements, suggesting that temporal information could benefit the prediction of diverse emergent behaviors. However, these improvement did not match the predictive accuracy of the final timepoints directly from the simulation. Based on a principal component analysis, the features derived from the forecasted network architectures were comparable to those from simulated architectures at the final timepoint, independent of the training-testing split of data (Supp. Figure 9). These results indicated that the propagation of minor errors resulting from ML regression techniques may account for the differences in predictive accuracy of the forecasted versus simulated timepoints.

## Discussion

Parity between biological systems and computational models requires algorithms capable of considering higher resolutions of heterogeneous species and their interactions. This objective adds complexity to analysis of experimental data and the development of predictive models. Emulation is a powerful tool that computational scientists can use to reduce the computational expense of model simulations (accelerating hypothesis generation), and to identify and understand key drivers of simulation dynamics (elucidating biological insight). This work unpacks the challenges of parity by using emulation to predict the evolution of tumor development in a dynamic environment that accounts for multilateral regulation among diverse cell agents and between cell agents and their supporting vasculature. The vascular architecture (topology) and function (hemodynamics) change as a function of time and cell population, encompassing a relevant level of biological complexity. While other studies have focused on leveraging emulation to interrogate ABM parameters (Alden et al., 2020; Angione et al., 2022), we focused on emulators using initial heterogeneous environmental conditions to predict the evolution of the cell population holistically to maintain close analogs with current experimental methodologies (e.g., dynamic imaging).

We built ML models designed to predict emergent behaviors of *in silico* tumors after two simulation weeks. These behaviors include tumor activity, growth, and symmetry. ML model predictions were based on features that derived from a network analysis of the tumor environment’s initial vascular architecture and hemodynamics. Counter-intuitively, we found that spatial characteristics of the environment were largely *un*successful at improving predictions of emergent behaviors that were shown to be associated with spatial phenomena (Yu and Bagheri, 2021). The resulting models were prone to both underfitting (evident from poor performance and limited improvement from additional data) and overfitting (indicated by substantial gaps between testing and training splits). Instead, we found that ML models that were trained on environmental snapshots of later simulation timepoints were more predictive of these emergent behaviors.

We then built an emulator using a combination of a RNN-forecaster model to predict the network evolution of the vasculature, and we used these as inputs into ML regression models to predict emergent behaviors. The final RNN-based emulator showed promising, albeit limited, improvement over using emulators that were strictly trained on initial environmental conditions highlighting the role of temporal dynamics in spatio-temporal cell population models. We were able to demonstrate the ability of an RNN model to make accurate forecasts of the network evolution. However, the limited predictive potential of those forecasted timepoints for predicting emergent behavior in turn emphasizes the importance of leveraging ABMs to make more accurate representations of our systems.

By leveraging network analysis and the evolution of network metrics, this study reinforces the importance of temporal information when predicting the behavior of cell population dynamics, even when the emergent behavior derives from spatial dynamics. The difference in performance between our two simulation contexts necessitate careful consideration when translating colony results to tissue contexts— analogous to translating in vitro to in vivo results. A general conclusion of this study suggests that out-of-the-box ML approaches are limited in their ability to characterize spatially heterogeneous systems. Despite their limited performance, we believe that SE would become an invaluable tool to the scientific community once we overcome challenges underlying their predictive performance for general application. Until then, ABMs are a necessary and unmatched alternative to predicting spatio-temporal dynamics.

The emulation of ABMs is difficult due to the complexity (e.g., number of species and corresponding interactions) and emergent nature of the biological phenomena they are well suited to simulate. As ABMs become increasingly multi-scale and complex (Prybutok et al., 2022a), challenges in emulation will persist and likely magnify. Mitigating these challenges is necessary to mediate the quantity of ABM simulations required to identify patterns, sample high-dimensional spaces, and generate testable hypotheses across emergent dynamics; such simulations can be cost-prohibitive. This cost is particularly relevant in context of personalized medicine where computational models must be used for real-time control (as is the case of insulin delivery)(Ortmann et al., 2019). In order to develop similar interventions that can help mitigate or drive population dynamics, the cost of predicting the spatio-temporal dynamics of cell populations must be addressed head on.

Computational expense remains a significant consideration in supporting emulator development, training, and analysis of perturbations’ impact on outcomes. Furthermore, the high-resolution spatio-temporal data required for emulation necessitates efficient storage, dissemination, and management protocols. The associated computational costs with emulation also call for economic considerations when investigating relationships among state variables or between inputs and outputs, as well as rigorous sampling of an ML model’s hyperparameter space. Additionally, emergent dynamics arising from the evolution of complex spatial and temporal processes poses challenges for representing said data in a ML framework such that the resulting models remain interpretable and useful. These challenges need to be addressed to maximize the potential of SE in advancing our understanding of biological systems through computational modeling.

## Acknowledgments

This work was supported by the National Science Foundation CAREER award CBET-1653315 (N.B.) and the Washington Research Foundation (N.B.).

## Emergent behavior metrics

Emergent behavior metrics are defined and calculated as presented in a previous study of heterogeneous vasculatures in ARCADE (Yu and Bagheri, 2021).

## Growth rate

Growth rate quantifies the change in colony diameter over time. First, colony diameter is calculated at each time index. Growth rate is the slope of the simple linear regression between [0, 0.5, . . ., *t_i_*] and corresponding diameters [*D*_0_, *D*_0.5_, . . ., *D_i_*], where *i* indicates the timepoint. These calculations were performed using the Python function polyfit from package numpy with degree of 1.

## Symmetry

Symmetry (*S*) quantifies colony shape at a given timepoint, ranging from 0 (not symmetric) to 1 (perfectly symmetric). For hexagonal coordinates, a colony is perfectly symmetric if for each location (u,v,w), the corresponding five locations (-w,-u,-v), (v,w,u), (-u,-v,-w), (w,u,v), and (-v,-w,-u) are all occupied. Symmetry is calculated as:

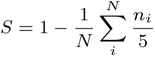

where *N* is the number of unique locations and *n_i_* is the number of corresponding unoccupied locations for a unique location *i*.

## Activity

Activity (*A*) quantifies the ratio of active (proliferative and migratory) to inactive (necrotic and apoptotic) cells at a given timepoint. This metric ranges between -1 (all non-quiescent cells are inactive) and +1 (all non-quiescent cells are active). An activity value of 0 indicates an equal number of active and inactive cells. Activity is calculated as:

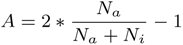

where *N_a_*is the number of active cells and *N_i_* is the number of inactive cells.

---

**Supp. Fig. 1:**
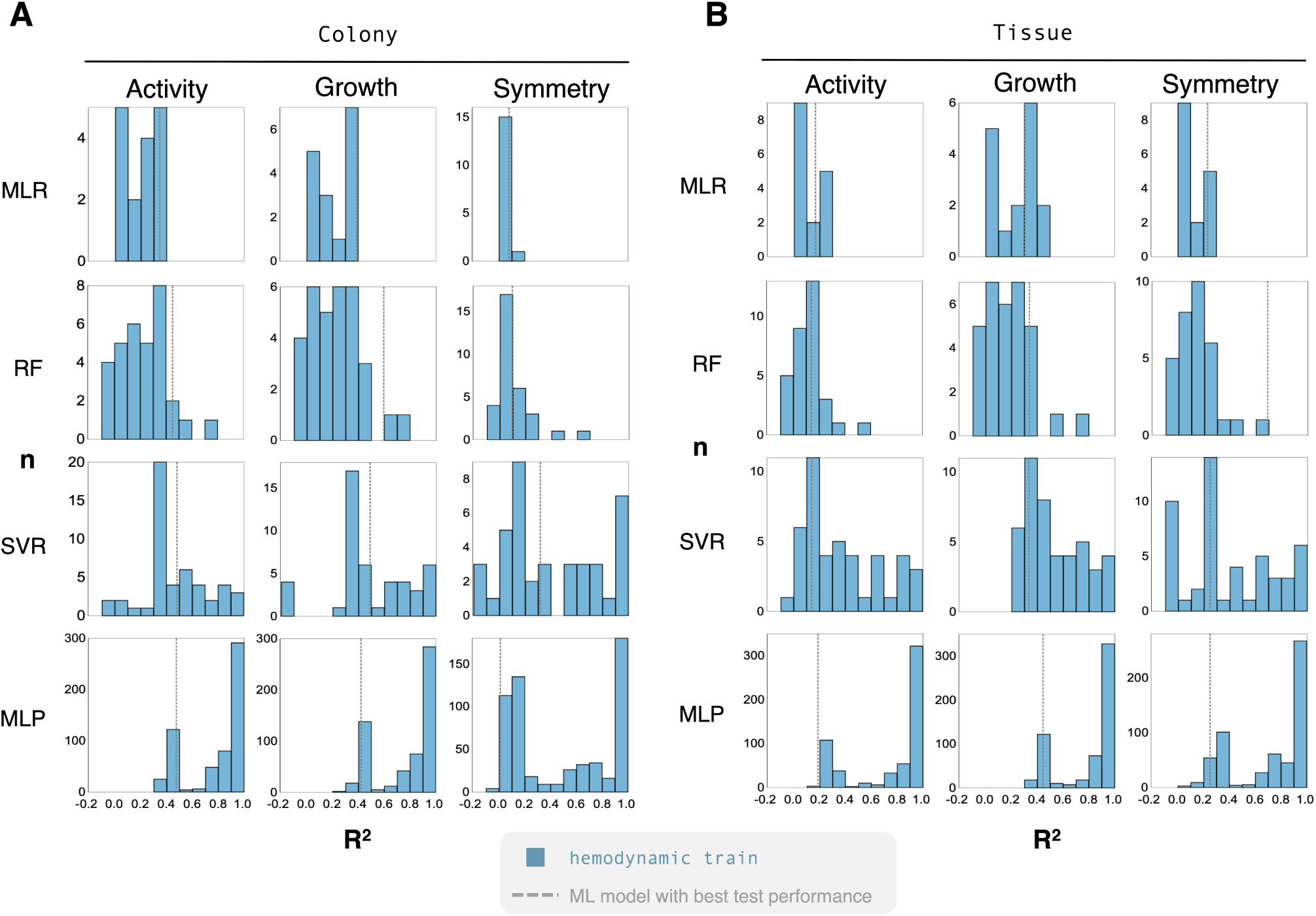
Emulators capture variance across the data during training. (**A**) Histograms show a range of performances achieved during training for different parameter combinations in the colony context. More complex models, such as MLP, can perfectly fit the data in many cases. The vertical dashed line indicates the training performance on model with the best validation performance. (**B**) Similarly, histograms show prediction performance during training across responses and models in the tissue context.

**Supp. Fig. 2:**
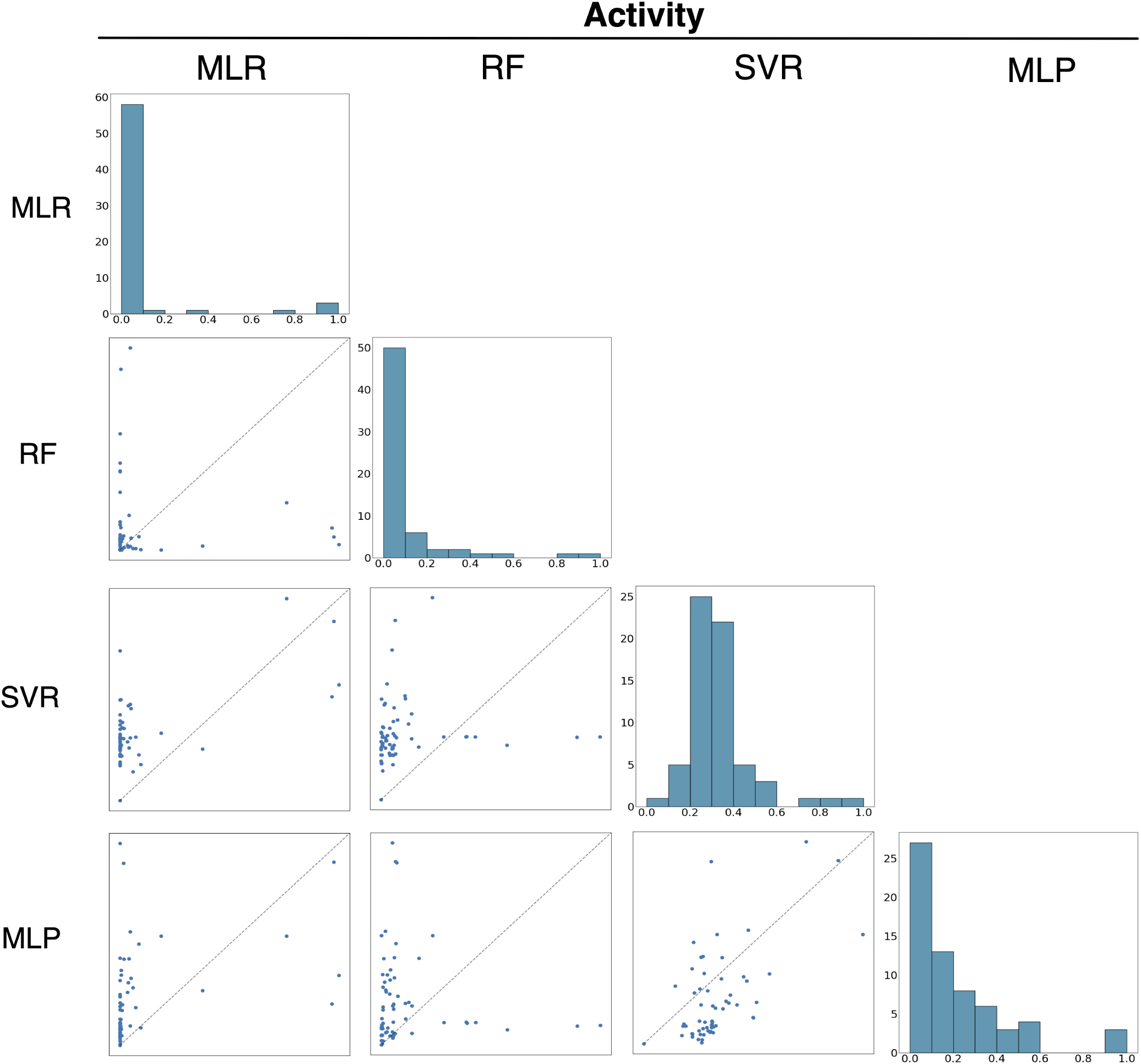
Discrepancy in feature importance across models. Histograms along the diagonal show the normalized importance distribution of features for each model. Parity plots show the importance of a feature for one model against another. Perfect parity would indicate that the models agreed on the importance level of each feature.

**Supp. Fig. 3:**
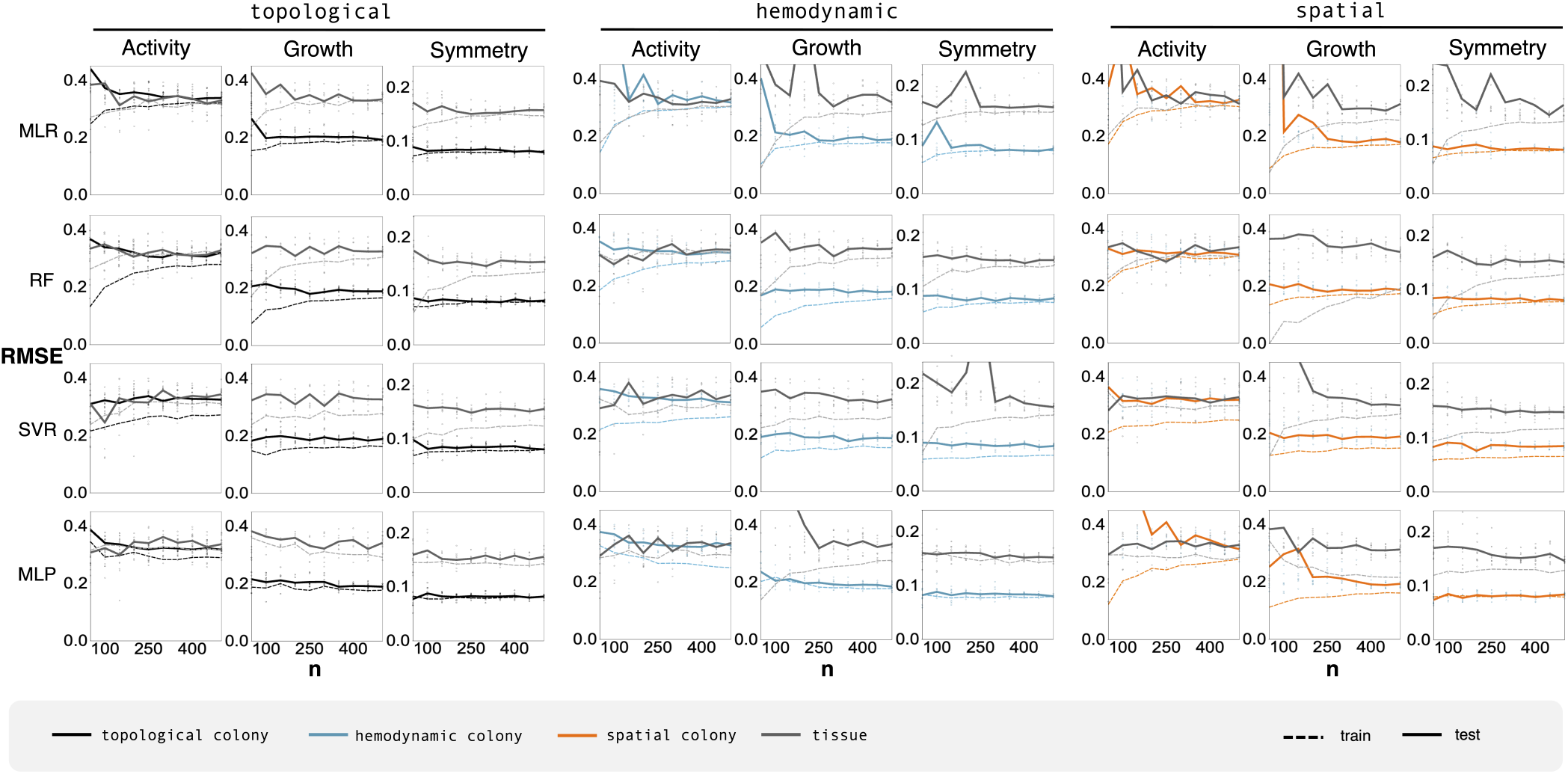
Training data points to diminishing returns. Line plots indicate the predictive performance of models trained on increasingly large training data sets. In most cases, the RMSE shows diminishing returns of model performance across all feature sets (topolgical, hemodynamic, and spatial), emergent targets (activity, growth, and symmetry), and algorithm (MRL, RF, SVR, MLP).

**Supp. Fig. 4:**
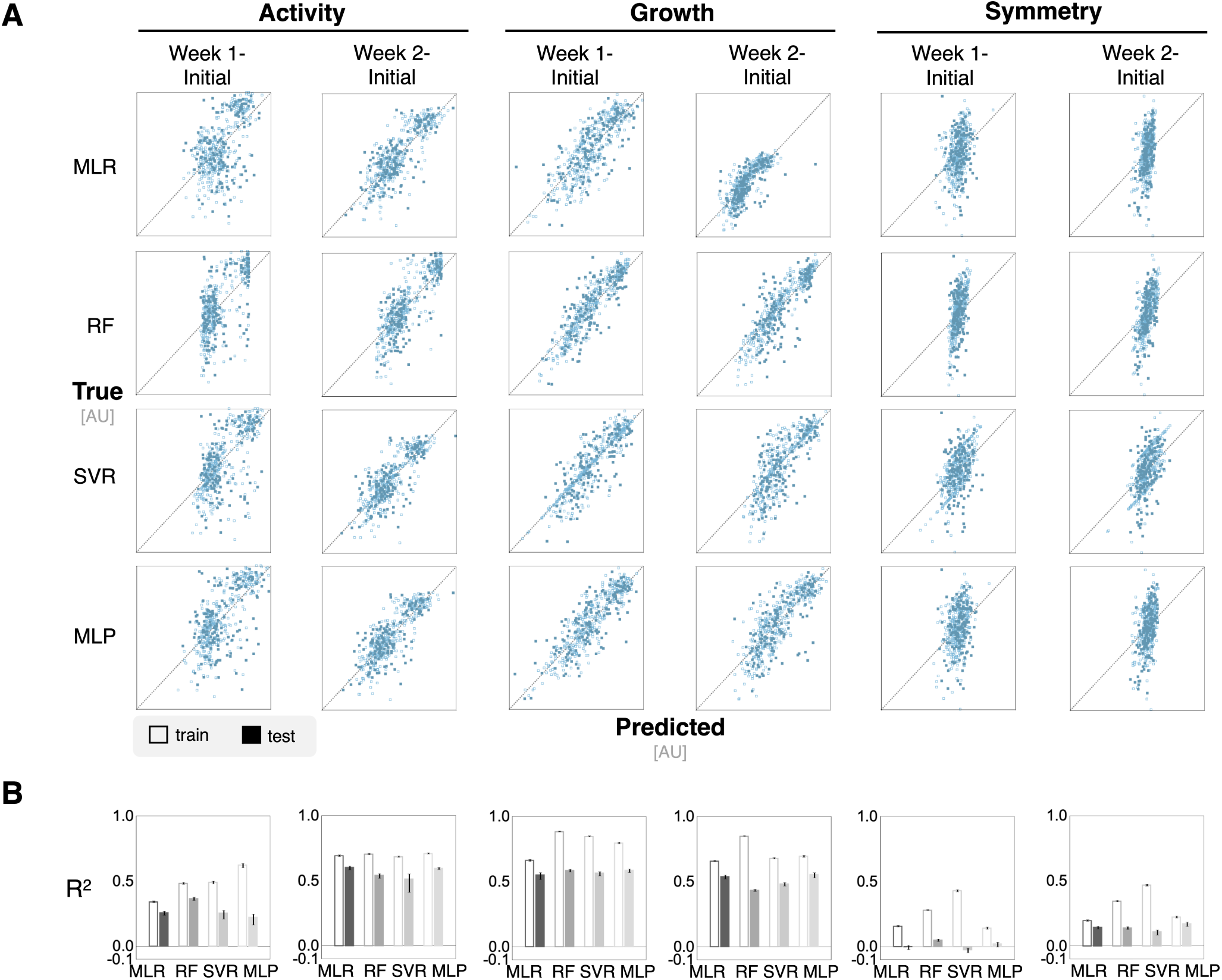
Differential timepoint analysis shows minimal improvement in performance in a colony context. Differential features were calculated by subtracting the features at either one or two weeks from initial features in order to capture the evolution of the features over time. (**A**) Parity plots show the predicted values of emergent targets against the real value to demonstrate fine-grained predictive performance. (**B**) Bar plots show *R*^2^ values for ML models trained on differential features. Growth predictions in the colony context improve with simulations using week one and initial timepoint differential features.

**Supp. Fig. 5:**
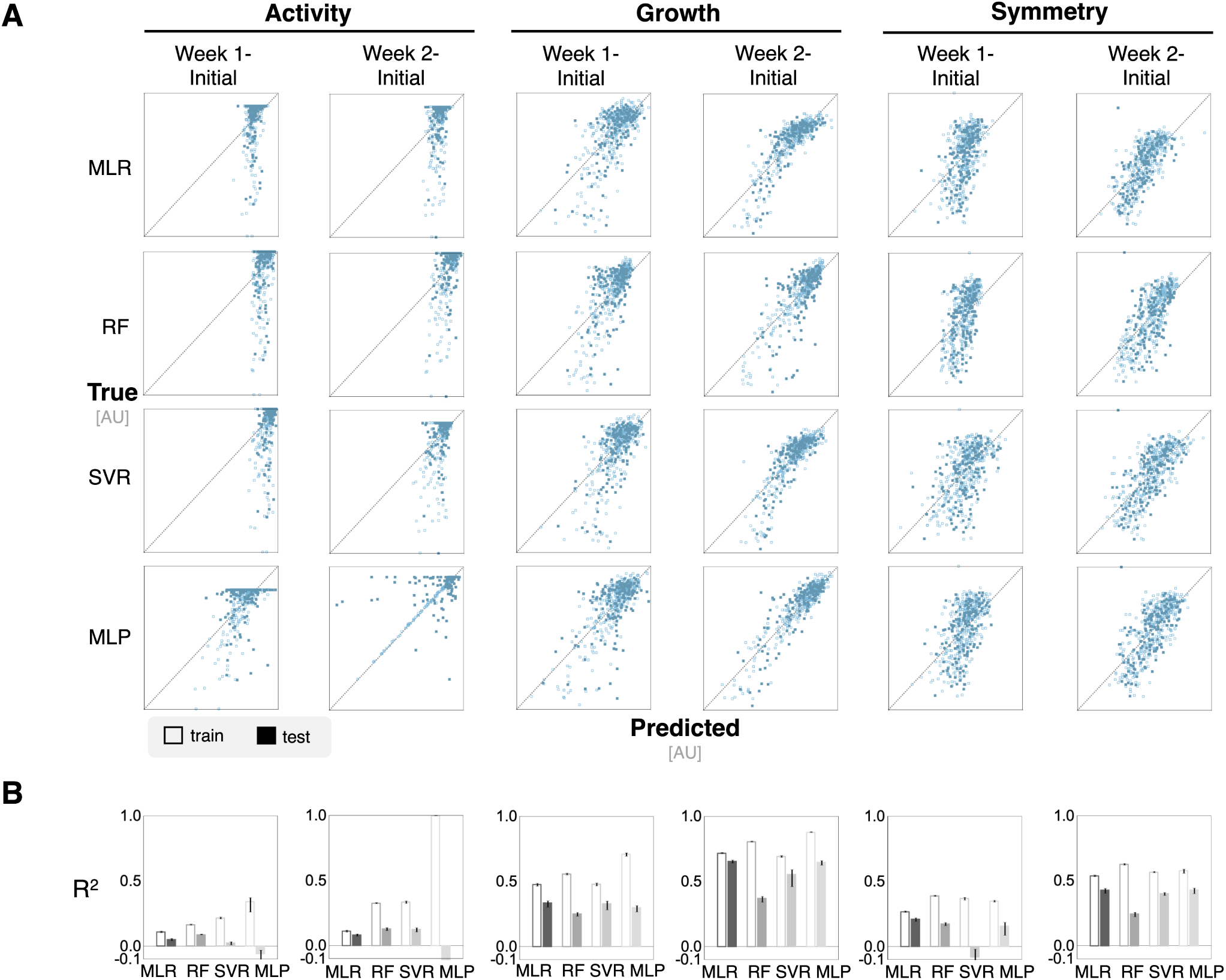
Differential timepoint analysis does not improve performance in a tissue context. Differential features were calculated by subtracting the features at either one or two weeks from initial features in order to capture the evolution of the features over time. (**A**) Parity plots show the predicted values of emergent targets against the real value to demonstrate fine-grained predictive performance. (**B**) Bar plots show *R*^2^ values for ML models trained on differential features.

**Supp. Fig. 6:**
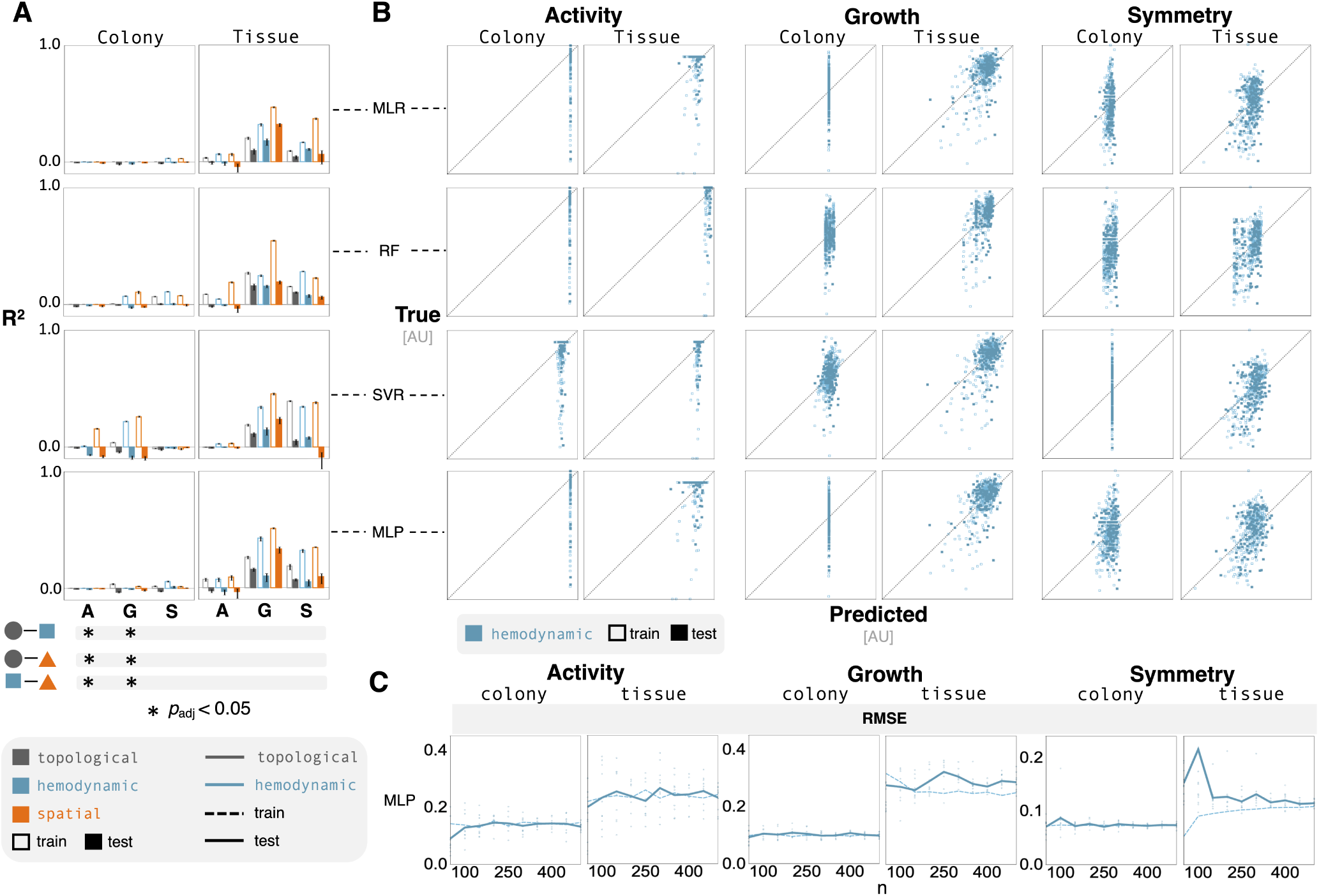
Shortened prediction horizon has minimal effect on emulator performance. Models predicting spatio-temporal dynamics from after one simulation week. (**A**) Bar plots indicate very poor predictive performance in all cases. Bar chart values range from -0.1 to 1.0; the horizontal axis is at 0.0. The Bonferroni corrected p-values from a two-way ANOVA highlight significant results (noted with black circles) that have an adjusted p-value less that 0.05. (**B**) Parity plots reveal substantial discrepancies in the variance between the predicted and true responses. (**C**) Line plots show predictive performance of the MLP models (the average RMSE) as a function of the size of training data. Performance improvements are limited from additional data. The data points highlight the RMSE from randomized test sets.

**Supp. Fig. 7:**
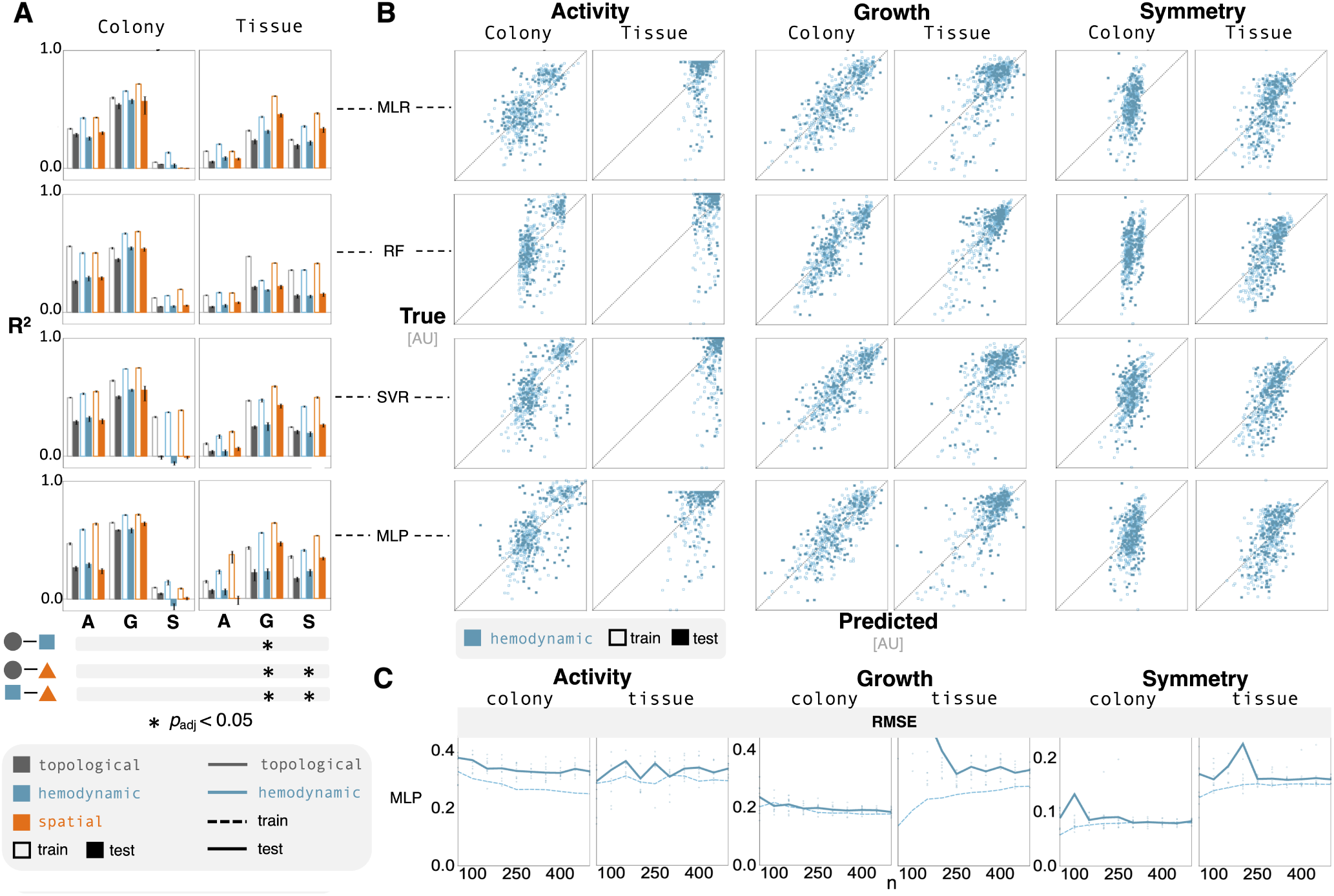
Mid-simulation features show improvement in performance in both contexts. Models trained on vascular features characterizing network structure at one simulation week result. (**A**) Bar plots show limited improvement from training on mid-simulation features. Bar chart values range from -0.1 to 1.0; the horizontal axis is at 0.0. The Bonferroni corrected p-values from a two-way ANOVA highlight significant results (noted with black circles) that have an adjusted p-value less that 0.05. (**B**) Parity plots reveal large amounts of variance in predicted values with some improvement in growth predictions in the colony context. (**C**) Line plots show predictive performance of the MLP models (the average RMSE) as a function of the size of training data. Performance improvements are limited from additional data. The data points highlight the RMSE from randomized test sets.

**Supp. Fig. 8:**
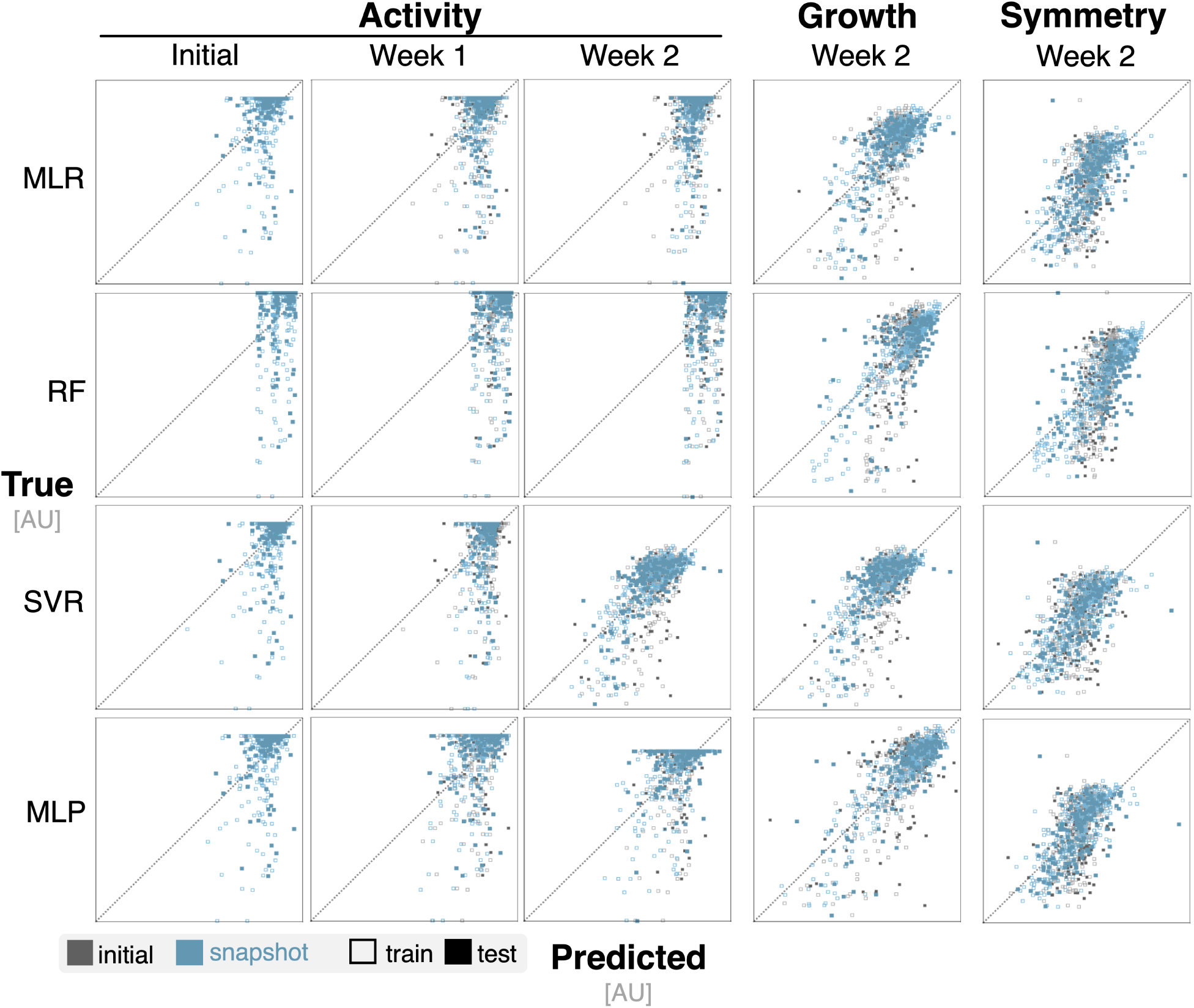
Temporal information improves ML model predictions in tissue context. Parity plots show the predictive performance of ML models trained on features from later timepoints against emulators trained on features from the initial timepoint. Improved prediction of activity is limited; growth and symmetry reflect minimal improvements.

**Supp. Fig. 9:**
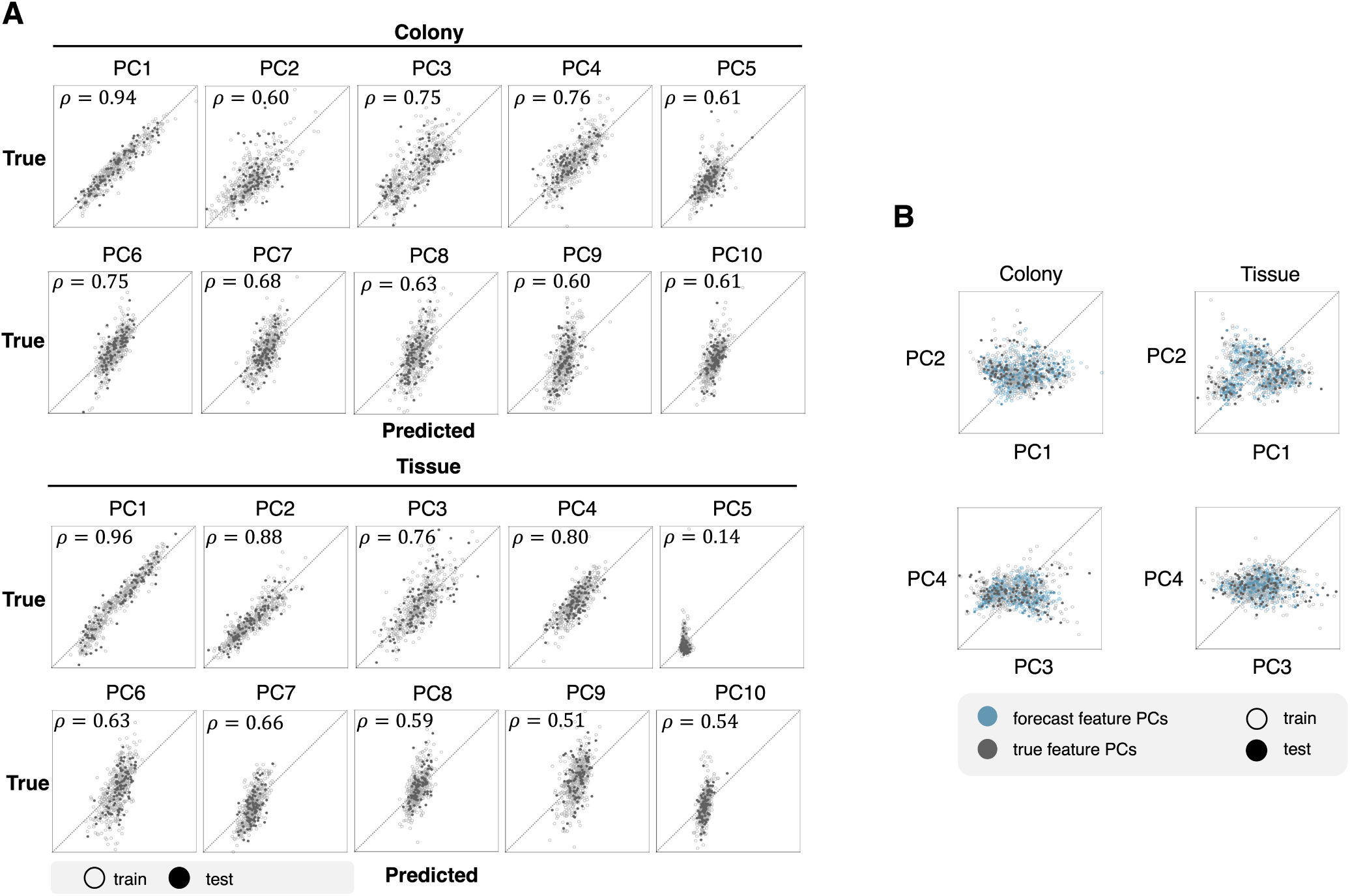
RNN model captures vascular feature variance. (**A**) Parity plots highlight the performance of a RNN model at predicting network metric features. Both the simulated features and forecasted features were combined to perform PCA. The dimensionality of the feature set are reduced to the first 10 principal components, which represent 95% of the feature variance. The Pearson correlation coefficient (*ρ*) is reported for each parity plot. (**B**) Scatter plots illustrate the overlap between true and forecasted features using the first four principal components.

**Supp. Fig. 10:**
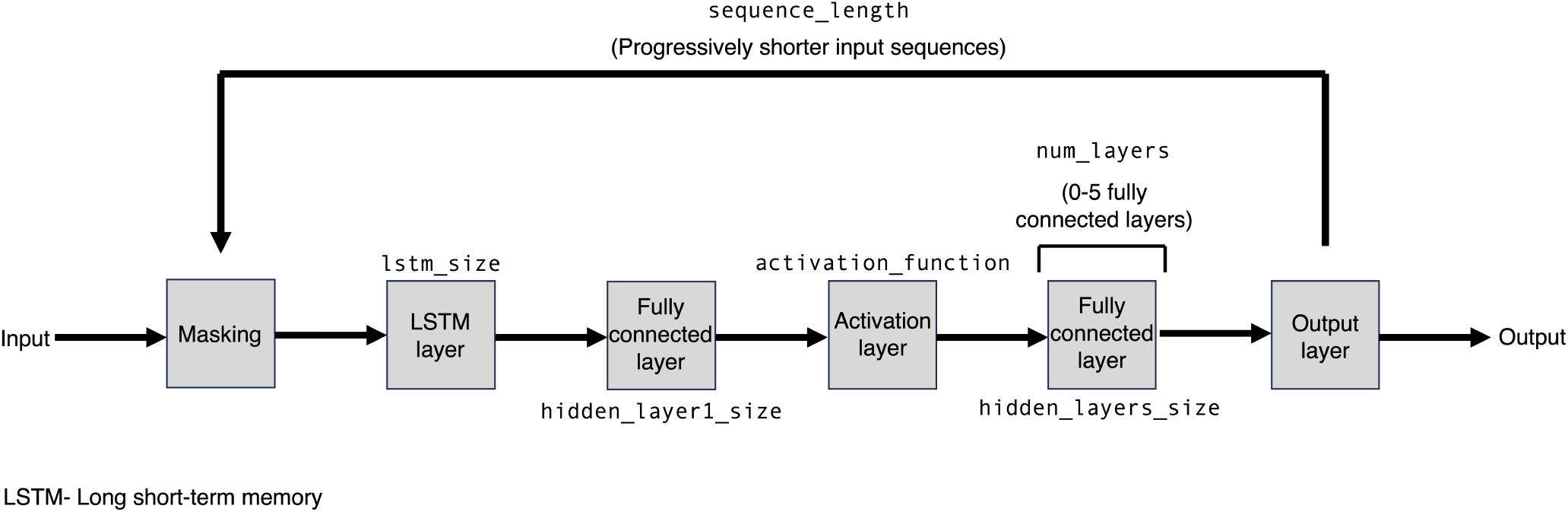
Architecture and structure of the RNN used for feature prediction. This flowchart describes the layers used while training the feature prediction RNN. All parameter values are defined in Supp. Table 7 and optimal values are provided in Supp. Table 8.

**Supp. Table 1.**
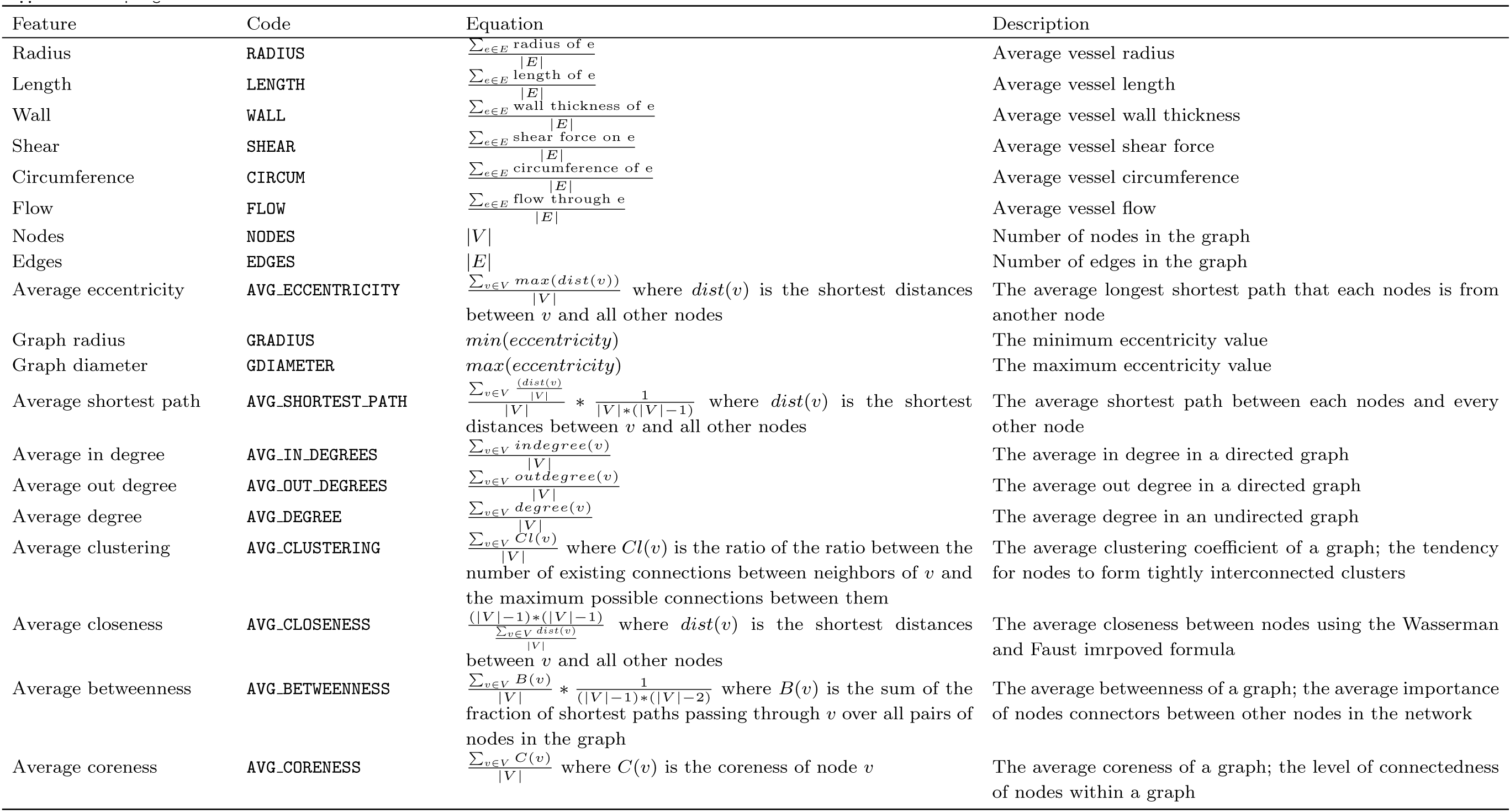
Topological feature list.

| Feature | Code | Equation | Description |
| --- | --- | --- | --- |
| Radius | RADIUS | $\sum_{e \in E} \frac{\text{radius of } e}{\vert E\vert }$ | Average vessel radius |
| Length | LENGTH | $\sum_{e \in E} \frac{\text{length of } e}{\vert E\vert }$ | Average vessel length |
| Wall | WALL | $\sum_{e \in E} \frac{\text{wall thickness of } e}{\vert E\vert }$ | Average vessel wall thickness |
| Shear | SHEAR | $\sum_{e \in E} \frac{\text{shear force on } e}{\vert E\vert }$ | Average vessel shear force |
| Circumference | CIRCUM | $\sum_{e \in E} \frac{\text{circumference of } e}{\vert E\vert }$ | Average vessel circumference |
| Flow | FLOW | $\sum_{e \in E} \frac{\text{flow through } e}{\vert E\vert }$ | Average vessel flow |
| Nodes | NODES | $\vert V\vert $ | Number of nodes in the graph |
| Edges | EDGES | $\vert E\vert $ | Number of edges in the graph |
| Average eccentricity | AVG_ECCENTRICITY | $\frac{\sum_{v \in V} \max(dist(v))}{\vert V\vert }$ where $dist(v)$ is the shortest distances between $v$ and all other nodes | The average longest shortest path that each nodes is from another node |
| Graph radius | GRADIUS | $\min(eccentricity)$ | The minimum eccentricity value |
| Graph diameter | GDIAMETER | $\max(eccentricity)$ | The maximum eccentricity value |
| Average shortest path | AVG_SHORTEST_PATH | $\frac{\sum_{v \in V} \frac{dist(v)}{\vert V\vert }}{\vert V\vert } * \frac{1}{\vert V\vert * (\vert V\vert - 1)}$ where $dist(v)$ is the shortest distances between $v$ and all other nodes | The average shortest path between each nodes and every other node |
| Average in degree | AVG_IN_DEGREES | $\sum_{v \in V} \frac{indegree(v)}{\vert V\vert }$ | The average in degree in a directed graph |
| Average out degree | AVG_OUT_DEGREES | $\sum_{v \in V} \frac{outdegree(v)}{\vert V\vert }$ | The average out degree in a directed graph |
| Average degree | AVG_DEGREE | $\sum_{v \in V} \frac{degree(v)}{\vert V\vert }$ | The average degree in an undirected graph |
| Average clustering | AVG_CLUSTERING | $\frac{\sum_{v \in V} Cl(v)}{\vert V\vert }$ where $Cl(v)$ is the ratio of the ratio between the number of existing connections between neighbors of $v$ and the maximum possible connections between them | The average clustering coefficient of a graph; the tendency for nodes to form tightly interconnected clusters |
| Average closeness | AVG_CLOSENESS | $\frac{(\vert V\vert - 1) * (\vert V\vert - 1)}{\sum_{v \in V} \frac{dist(v)}{\vert V\vert }}$ where $dist(v)$ is the shortest distances between $v$ and all other nodes | The average closeness between nodes using the Wasserman and Faust improved formula |
| Average betweenness | AVG_BETWEENNESS | $\frac{\sum_{v \in V} B(v)}{\vert V\vert } * \frac{1}{(\vert V\vert - 1) * (\vert V\vert - 2)}$ where $B(v)$ is the sum of the fraction of shortest paths passing through $v$ over all pairs of nodes in the graph | The average betweenness of a graph; the average importance of nodes connectors between other nodes in the network |
| Average coreness | AVG_CORENESS | $\frac{\sum_{v \in V} C(v)}{\vert V\vert }$ where $C(v)$ is the coreness of node $v$ | The average coreness of a graph; the level of connectedness of nodes within a graph |

**Supp. Table 2.**
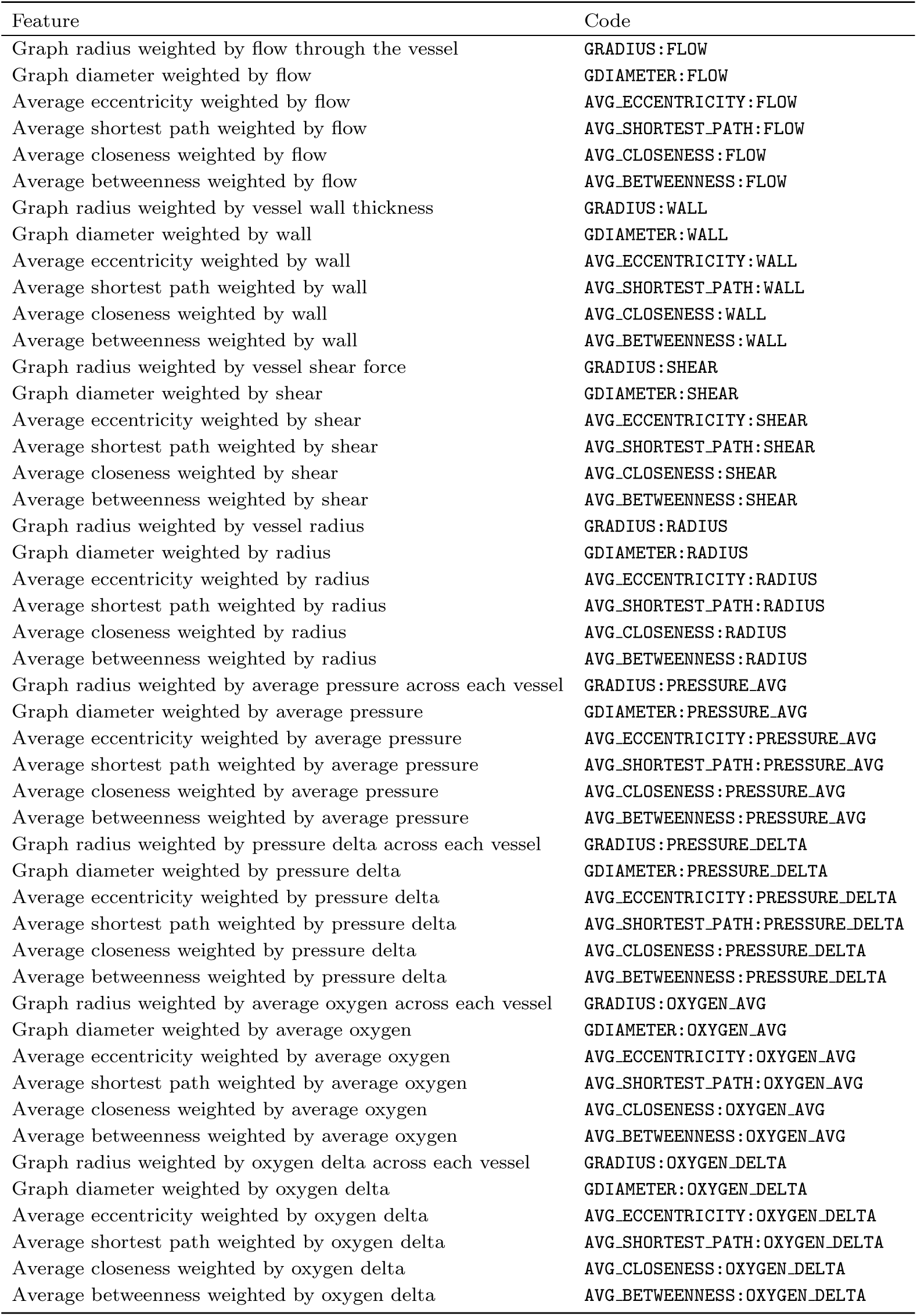
Hemodynamic feature list.

**Supp. Table 3.**
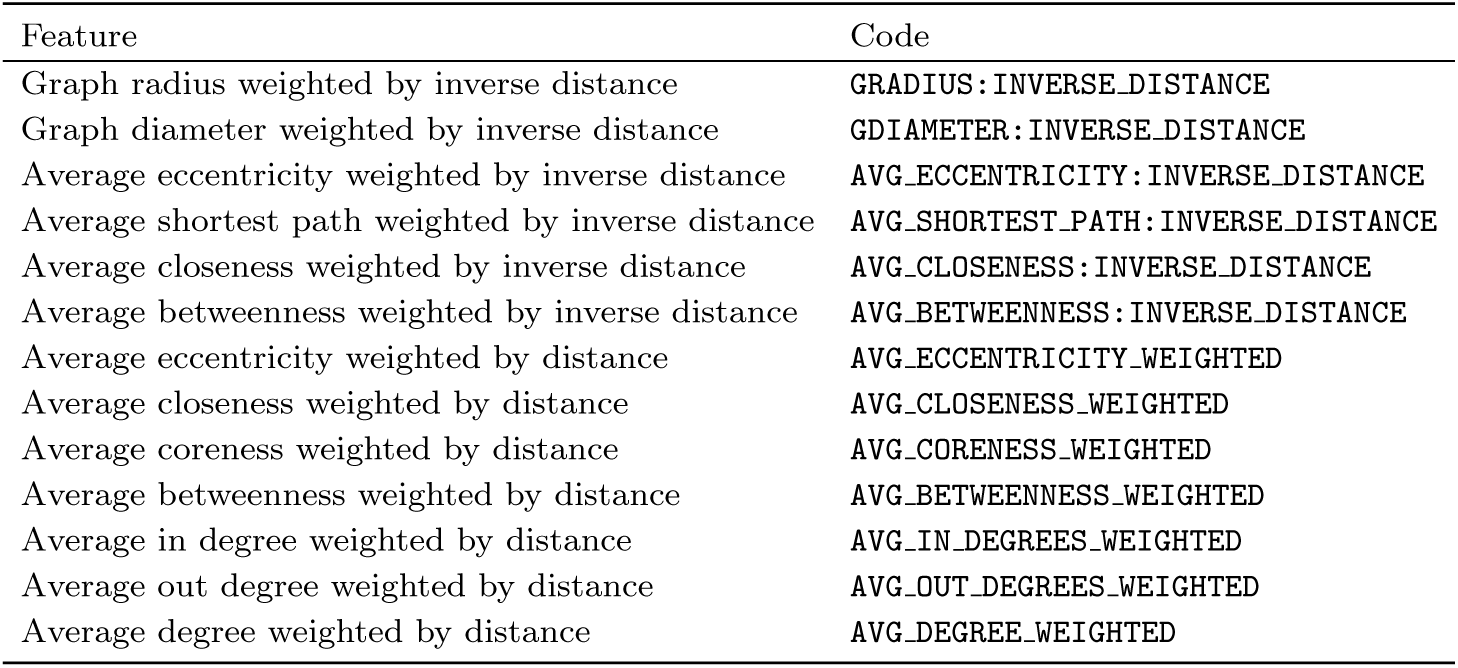
Spatial feature list.

**Supp. Table 4.**
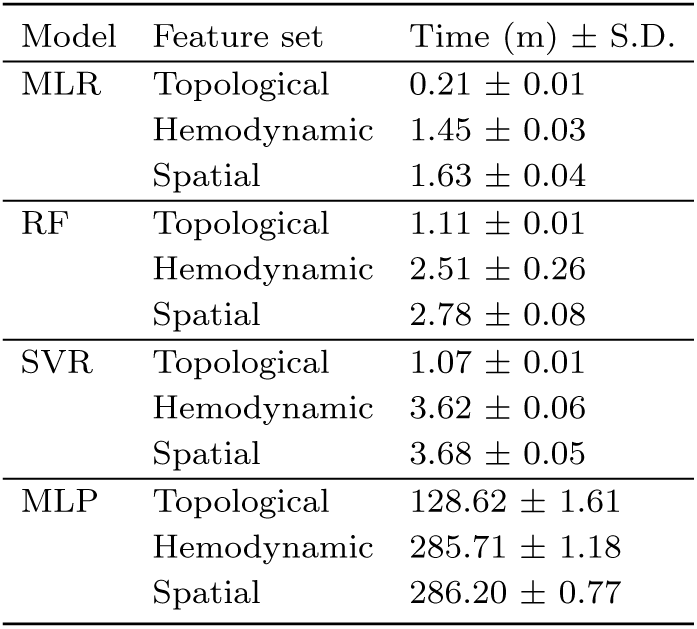
Emulator run times across feature sets on an m5.large EC2 instance.

| Model | Feature set | Time (m) ± S.D. |
| --- | --- | --- |
| MLR | Topological | 0.21 ± 0.01 |
| | Hemodynamic | 1.45 ± 0.03 |
| | Spatial | 1.63 ± 0.04 |
| RF | Topological | 1.11 ± 0.01 |
| | Hemodynamic | 2.51 ± 0.26 |
| | Spatial | 2.78 ± 0.08 |
| SVR | Topological | 1.07 ± 0.01 |
| | Hemodynamic | 3.62 ± 0.06 |
| | Spatial | 3.68 ± 0.05 |
| MLP | Topological | 128.62 ± 1.61 |
| | Hemodynamic | 285.71 ± 1.18 |
| | Spatial | 286.20 ± 0.77 |

**Supp. Table 5.**
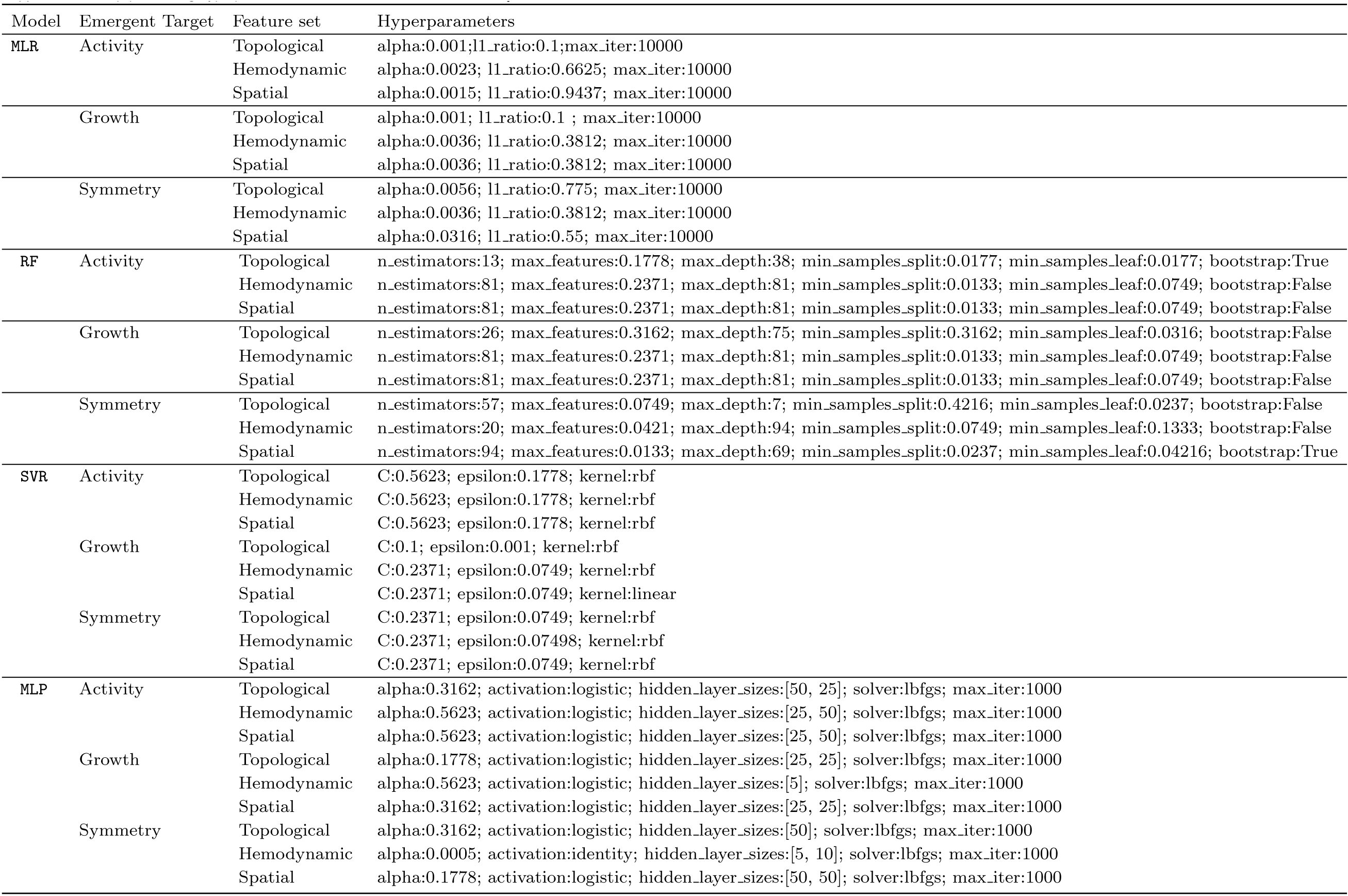
Top performing hyperparameters for emulation models in a colony context.

**Supp. Table 6.**
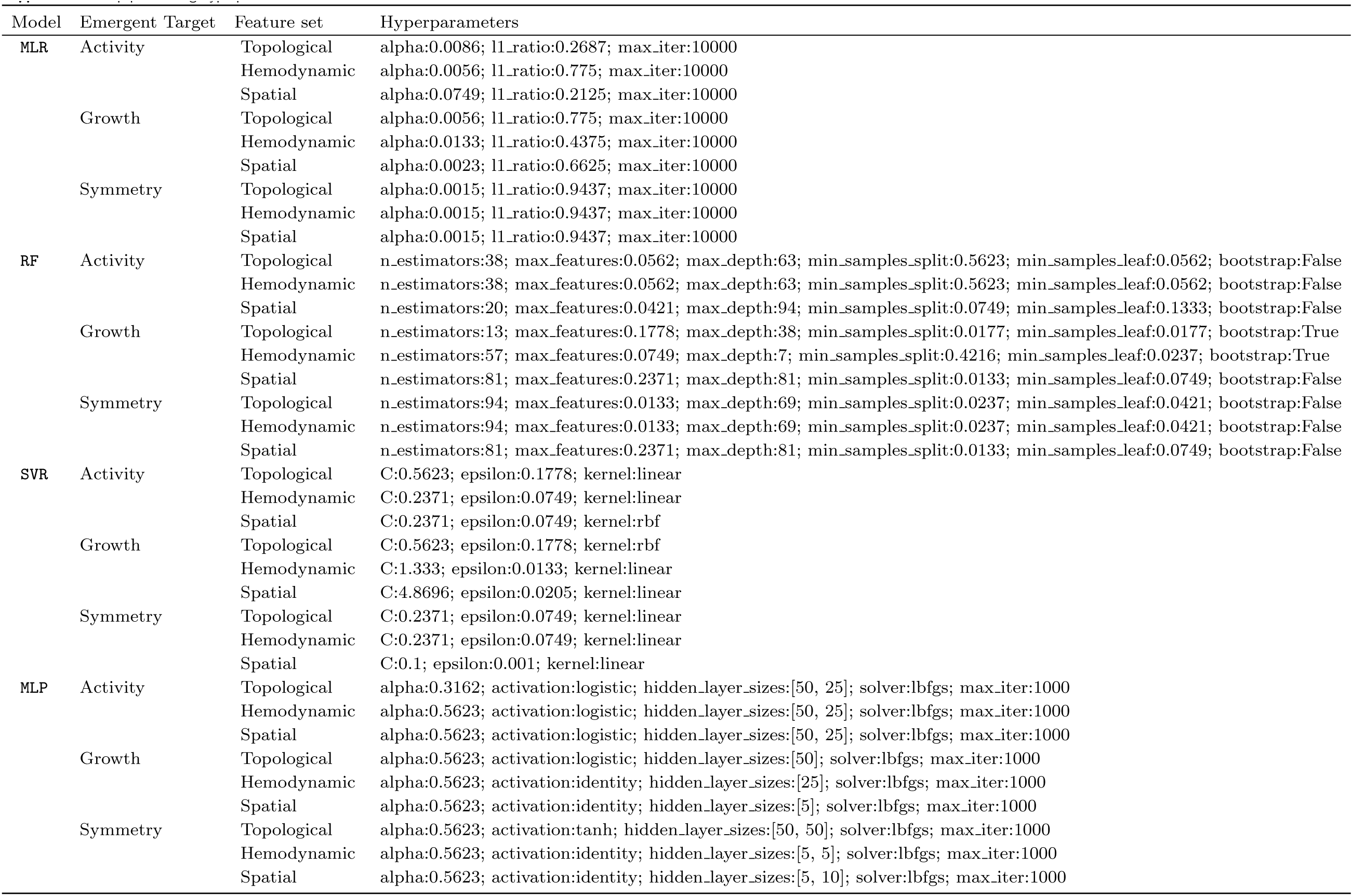
Top performing hyperparameters for emulation models in a tissue context.

**Supp. Table 7.**
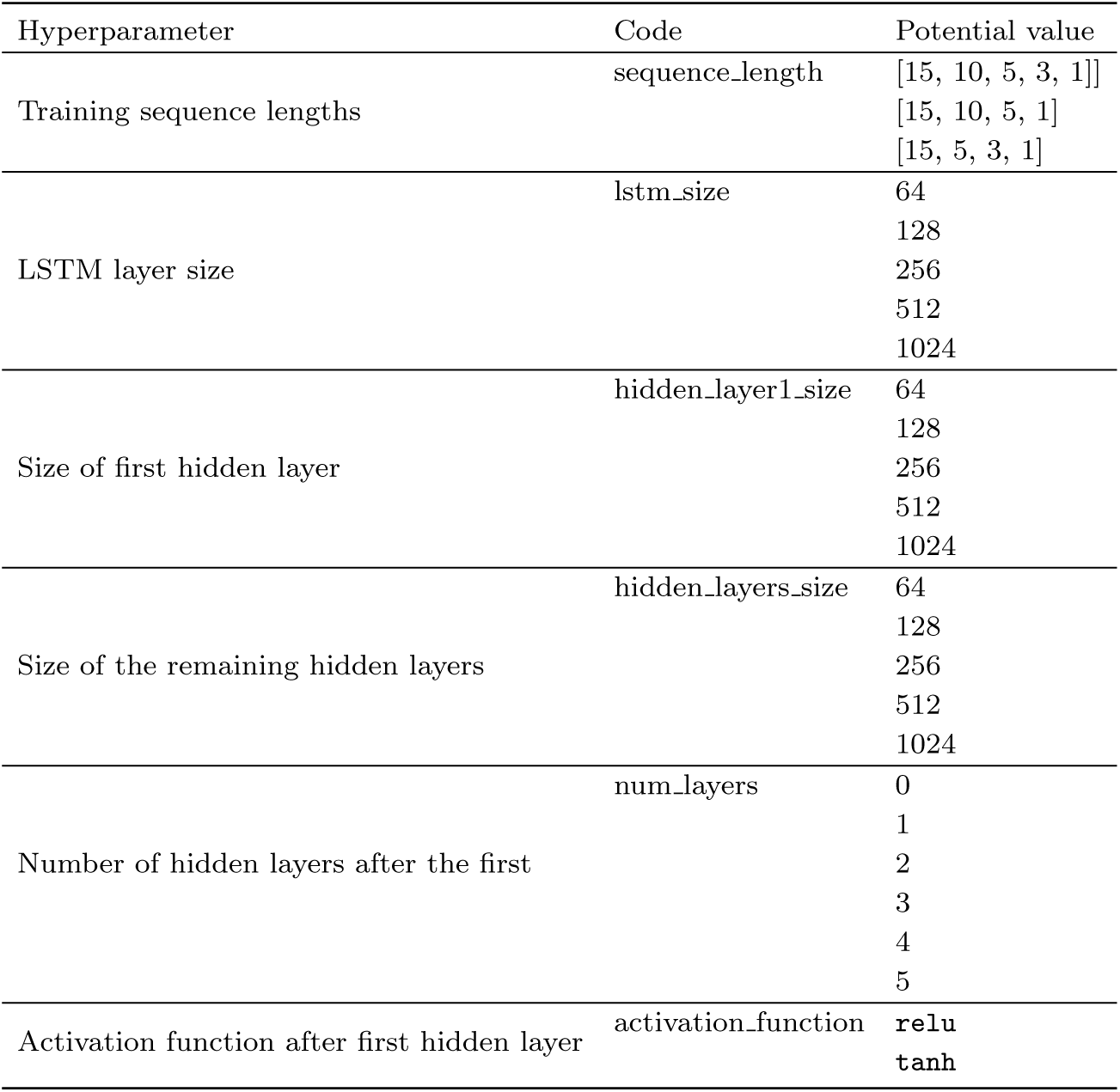
Hyperparameters used in RNN architecture grid search.

**Supp. Table 8.**
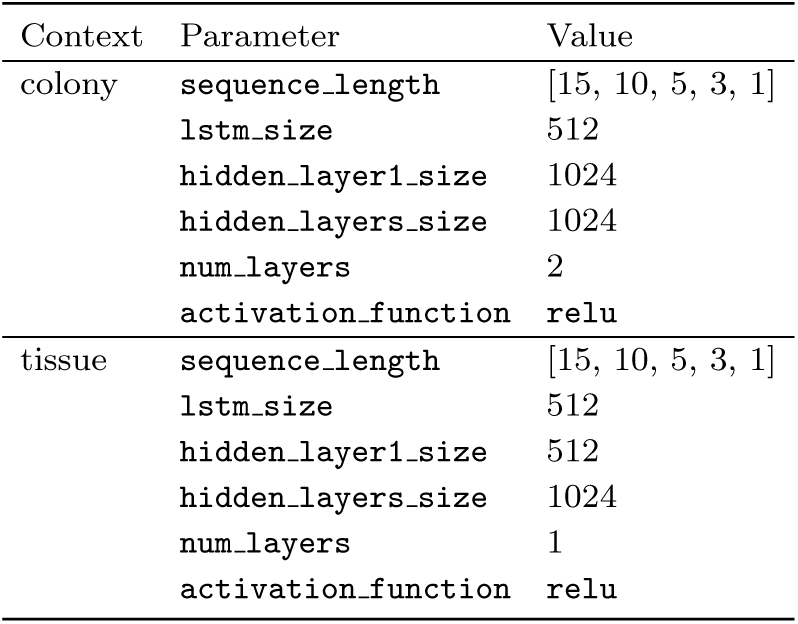
Top performing architecture parameters for RNN.

## References

1. M. Abadi, A. Agarwal, P. Barham, E. Brevdo, Z. Chen, C. Citro, G. S. Corrado, A. Davis, J. Dean, M. Devin, S. Ghemawat, I. Goodfellow, A. Harp, G. Irving, M. Isard, Y. Jia, R. Jozefowicz, L. Kaiser, M. Kudlur, J. Levenberg, D. Mané, R. Monga, S. Moore, D. Murray, C. Olah, M. Schuster, J. Shlens, B. Steiner, I. Sutskever, K. Talwar, P. Tucker, V. Vanhoucke, V. Vasudevan, F. Viégas, O. Vinyals, P. Warden, M. Wattenberg, M. Wicke, Y. Yu, and X. Zheng. TensorFlow: Large-scale machine learning on heterogeneous systems, 2015. URL https://www.tensorflow.org/. Software available from tensorflow.org.

2. K. Alden, J. Cosgrove, M. Coles, and J. Timmis. Using Emulation to Engineer and Understand Simulations of Biological Systems. IEEE/ACM Trans Comput Biol Bioinform, 17(1):302–315, 2020. ISSN 1557-9964. doi: 10.1109/TCBB.2018.2843339.

3. A. P. Alves, O. N. Mesquita, J. Gómez-Gardeñes, and U. Agero. Graph analysis of cell clusters forming vascular networks. R Soc Open Sci, 5(3), Mar. 2018. ISSN 2054-5703. doi: 10.1098/rsos.171592.

4. I. Amat-Roldan, A. Berzigotti, R. Gilabert, and J. Bosch. Assessment of Hepatic Vascular Network Connectivity with Automated Graph Analysis of Dynamic Contrast-enhanced US to Evaluate Portal Hypertension in Patients with Cirrhosis: A Pilot Study. Radiology, 277(1):268–276, Oct. 2015. ISSN 0033-8419. doi: 10.1148/radiol.2015141941.

5. C. Angione, E. Silverman, and E. Yaneske. Using machine learning as a surrogate model for agent-based simulations. PLOS ONE, 17 (2):e0263150, Feb. 2022. ISSN 1932-6203. doi: 10.1371/journal.pone.0263150.

6. N. Bagheri, A. E. Carpenter, E. Lundberg, A. L. Plant, and R. Horwitz. The new era of quantitative cell imaging—challenges and opportunities. Molecular Cell, 82(2):241–247, Jan. 2022. ISSN 1097-2765. doi: 10.1016/j.molcel.2021.12.024.

7. D. S. Bassett and O. Sporns. Network neuroscience. Nat Neurosci, 20(3):353–364, Mar. 2017. ISSN 1546-1726. doi: 10.1038/nn.4502.

8. G. Bindea, B. Mlecnik, M. Tosolini, A. Kirilovsky, M. Waldner, A. C. Obenauf, H. Angell, T. Fredriksen, L. Lafontaine, A. Berger, P. Bruneval, W. H. Fridman, C. Becker, F. Pages, M. R. Speicher, Z. Trajanoski, and J. Galon. Spatiotemporal Dynamics of Intratumoral Immune Cells Reveal the Immune Landscape in Human Cancer. Immunity, 39(4):782–795, Oct. 2013. ISSN 1074-7613. doi: 10.1016/j.immuni.2013.10.003.

9. P. J. Blanco, L. O. Müller, and J. D. Spence. Blood pressure gradients in cerebral arteries: A clue to pathogenesis of cerebral small vessel disease. Stroke Vasc Neurol, 2(3), Sept. 2017. ISSN 2059-8688, 2059-8696. doi: 10.1136/svn-2017-000087.

10. E. Bonabeau. Agent-based modeling: Methods and techniques for simulating human systems. Proceedings of the National Academy of Sciences, 99(suppl 3):7280–7287, May 2002. doi: 10.1073/pnas.082080899.

11. C. G. Cess and S. D. Finley. Multi-scale modeling of macrophage—T cell interactions within the tumor microenvironment. PLOS Computational Biology, 16(12):e1008519, Dec. 2020. ISSN 1553-7358. doi: 10.1371/journal.pcbi.1008519.

12. F. Chollet et al. Keras, 2015. URL https://github.com/fchollet/keras.

13. J. M. Cicchese, E. Pienaar, D. E. Kirschner, and J. J. Linderman. Applying optimization algorithms to tuberculosis antibiotic treatment regimens. Cellular and molecular bioengineering, 10:523–535, 2017.

14. A. Corti, M. Colombo, F. Migliavacca, J. F. Rodriguez Matas, S. Casarin, and C. Chiastra. Multiscale Computational Modeling of Vascular Adaptation: A Systems Biology Approach Using Agent-Based Models. Frontiers in Bioengineering and Biotechnology, 9, 2021. ISSN 2296-4185.

15. G. Csardi and T. Nepusz. The Igraph Software Package for Complex Network Research. *InterJournal*, Complex Systems:1695, Nov. 2005.

16. C. W. Dunnett. A multiple comparison procedure for comparing several treatments with a control. Journal of the American Statistical Association, 50(272):1096–1121, 1955. doi: 10.1080/01621459.1955.10501294. URL https://www.tandfonline.com/doi/abs/10.1080/01621459.1955.10501294.

17. R. Eftimie. Grand challenges in mathematical biology: Integrating multi-scale modeling and data. Frontiers in Applied Mathematics and Statistics, 8, 2022. ISSN 2297-4687.

18. R. Eftimie, A. Mavrodin, and S. P. A. Bordas. Chapter Four - From digital control to digital twins in medicine: A brief review and future perspectives. In S. P. A. Bordas, editor, Advances in Applied Mechanics, volume 56, pages 323–368. Elsevier, Jan. 2023. doi: 10.1016/bs.aams.2022.09.001.

19. A. Fouladzadeh, M. Dorraki, K. K. M. Min, M. P. Cockshell, E. J. Thompson, J. W. Verjans, A. Allison, C. S. Bonder, and D. Abbott. The development of tumour vascular networks. Commun Biol, 4(1):1–10, Sept. 2021. ISSN 2399-3642. doi: 10.1038/s42003-021-02632-x.

20. T. Fredrich, M. Welter, and H. Rieger. Tumorcode. Eur. Phys. J. E, 41(4):55, Apr. 2018. ISSN 1292-895X. doi: 10.1140/epje/i2018-11659-x.

21. T. Fredrich, H. Rieger, R. Chignola, and E. Milotti. Fine-grained simulations of the microenvironment of vascularized tumours. Sci Rep, 9(1):11698, Aug. 2019. ISSN 2045-2322. doi: 10.1038/s41598-019-48252-8.

22. A. Ghaffarizadeh, R. Heiland, S. H. Friedman, S. M. Mumenthaler, and P. Macklin. PhysiCell: An open source physics-based cell simulator for 3-D multicellular systems. PLOS Computational Biology, 14(2):e1005991, Feb. 2018. ISSN 1553-7358. doi: 10.1371/journal.pcbi.1005991.

23. C. M. Glen, M. L. Kemp, and E. O. Voit. Agent-based modeling of morphogenetic systems: Advantages and challenges. PLOS Computational Biology, 15(3):e1006577, Mar. 2019. ISSN 1553-7358. doi:10.1371/journal.pcbi.1006577.

24. J. Gomez, J. Prieto, E. Leon, and A. Rodríguez. INFEKTA—An agent-based model for transmission of infectious diseases: The COVID-19 case in Bogotá, Colombia. PLoS One, 16(2):e0245787, Feb. 2021. ISSN 1932-6203. doi: 10.1371/journal.pone.0245787.

25. D. Hanahan and R. A. Weinberg. Hallmarks of Cancer: The Next Generation. Cell, 144(5):646–674, Mar. 2011. ISSN 0092-8674. doi: 10.1016/j.cell.2011.02.013.

26. A. Heppenstall, A. Crooks, N. Malleson, E. Manley, J. Ge, and M. Batty. Future Developments in Geographical Agent-Based Models: Challenges and Opportunities. Geographical Analysis, 53(1):76–91, 2021. ISSN 1538-4632. doi: 10.1111/gean.12267.

27. L. Heumos, A. C. Schaar, C. Lance, A. Litinetskaya, F. Drost, L. Zappia, M. D. Lücken, D. C. Strobl, J. Henao, F. Curion, H. B. Schiller, and F. J. Theis. Best practices for single-cell analysis across modalities. Nat Rev Genet, pages 1–23, Mar. 2023. ISSN 1471-0064. doi: 10.1038/s41576-023-00586-w.

28. S. Hochreiter and J. Schmidhuber. Long short-term memory. Neural computation, 9(8):1735–1780, 1997.

29. E. Hunter, B. M. Namee, and J. Kelleher. An open-data-driven agent-based model to simulate infectious disease outbreaks. PLOS ONE, 13(12):e0208775, Dec. 2018. ISSN 1932-6203. doi: 10.1371/journal.pone.0208775.

30. B. Hwang, J. H. Lee, and D. Bang. Single-cell RNA sequencing technologies and bioinformatics pipelines. Exp Mol Med, 50(8):1–14, Aug. 2018. ISSN 2092-6413. doi: 10.1038/s12276-018-0071-8.

31. S. Iadevaia, Y. Lu, F. C. Morales, G. B. Mills, and P. T. Ram. Identification of Optimal Drug Combinations Targeting Cellular Networks: Integrating Phospho-Proteomics and Computational Network Analysis. Cancer Research, 70(17):6704–6714, Aug. 2010. ISSN 0008-5472. doi: 10.1158/0008-5472.CAN-10-0460.

32. Z. Ji, K. Yan, W. Li, H. Hu, and X. Zhu. Mathematical and Computational Modeling in Complex Biological Systems. BioMed Research International, 2017:e5958321, Mar. 2017. ISSN 2314-6133. doi: 10.1155/2017/5958321.

33. M. E. Johnson, A. Chen, J. R. Faeder, P. Henning, I. I. Moraru, M. Meier-Schellersheim, R. F. Murphy, T. Prüstel, J. A. Theriot, and A. M. Uhrmacher. Quantifying the roles of space and stochasticity in computer simulations for cell biology and cellular biochemistry. MBoC, 32(2):186–210, Jan. 2021. ISSN 1059-1524. doi: 10.1091/mbc.E20-08-0530.

34. C. C. Kerr, R. M. Stuart, D. Mistry, R. G. Abeysuriya, K. Rosenfeld, G. R. Hart, R. C. Núñez, J. A. Cohen, P. Selvaraj, B. Hagedorn, L. George, M. Jastrzebski, A. S. Izzo, G. Fowler, A. Palmer, D. Delport, N. Scott, S. L. Kelly, C. S. Bennette, B. G. Wagner, S. T. Chang, A. P. Oron, E. A. Wenger, J. Panovska-Griffiths, M. Famulare, and D. J. Klein. Covasim: An agent-based model of COVID-19 dynamics and interventions. PLOS Computational Biology, 17(7):e1009149, July 2021. ISSN 1553-7358. doi: 10.1371/journal.pcbi.1009149.

35. M. Kieu, H. Nguyen, J. A. Ward, and N. Malleson. Towards real-time predictions using emulators of agent-based models. Journal of Simulation, 0(0):1–18, June 2022. ISSN 1747-7778. doi: 10.1080/17477778.2022.2080008.

36. J. G. Kok, A. Leemans, L. K. Teune, K. L. Leenders, M. J. McKeown, S. Appel-Cresswell, H. P. H. Kremer, and B. M. de Jong. Structural Network Analysis Using Diffusion MRI Tractography in Parkinson’s Disease and Correlations With Motor Impairment. Front Neurol, 11:841, Sept. 2020. ISSN 1664-2295. doi: 10.3389/fneur.2020.00841.

37. M. Koutrouli, E. Karatzas, D. Paez-Espino, and G. A. Pavlopoulos. A Guide to Conquer the Biological Network Era Using Graph Theory. Frontiers in Bioengineering and Biotechnology, 8, 2020. ISSN 2296-4185.

38. D. Lähnemann, J. Köster, E. Szczurek, D. J. McCarthy, S. C. Hicks, M. D. Robinson, C. A. Vallejos, K. R. Campbell, N. Beerenwinkel, A. Mahfouz, L. Pinello, P. Skums, A. Stamatakis, C. S.-O. Attolini, S. Aparicio, J. Baaijens, M. Balvert, B. de Barbanson, A. Cappuccio, G. Corleone, B. E. Dutilh, M. Florescu, V. Guryev, R. Holmer, K. Jahn, T. J. Lobo, E. M. Keizer, I. Khatri, S. M. Kielbasa, J. O. Korbel, A. M. Kozlov, T.-H. Kuo, B. P. Lelieveldt, I. I. Mandoiu, J. C. Marioni, T. Marschall, F. Mölder, A. Niknejad, L. Raczkowski, M. Reinders, J. de Ridder, A.-E. Saliba, A. Somarakis, O. Stegle, F. J. Theis, H. Yang, A. Zelikovsky, A. C. McHardy, B. J. Raphael, S. P. Shah, and A. Schönhuth. Eleven grand challenges in single-cell data science. Genome Biology, 21(1):31, Feb. 2020. ISSN 1474-760X. doi: 10.1186/s13059-020-1926-6.

39. M. Lelek, M. T. Gyparaki, G. Beliu, F. Schueder, J. Griffíe, S. Manley, R. Jungmann, M. Sauer, M. Lakadamyali, and C. Zimmer. Single-molecule localization microscopy. Nat Rev Methods Primers, 1:39, 2021. ISSN 2662-8449. doi: 10.1038/s43586-021-00038-x.

40. C. S. Magnano and A. Gitter. Automating parameter selection to avoid implausible biological pathway models. npj Syst Biol Appl, 7 (1):1–12, Feb. 2021. ISSN 2056-7189. doi: 10.1038/s41540-020-00167-1.

41. J. Metzcar, Y. Wang, R. Heiland, and P. Macklin. A review of cell-based computational modeling in cancer biology. JCO Clinical Cancer Informatics, (3):1–13, 2019. doi: 10.1200/CCI.18.00069. URL https://doi.org/10.1200/CCI.18.00069. PMID: 30715927.

42. G. Modica, S. Praticò, L. Laudari, A. Ledda, S. Di Fazio, and A. De Montis. Implementation of multispecies ecological networks at the regional scale: Analysis and multi-temporal assessment. Journal of Environmental Management, 289:112494, July 2021. ISSN 0301-4797. doi: 10.1016/j.jenvman.2021.112494.

43. J. Möller and R. Pörtner. Digital Twins for Tissue Culture Techniques—Concepts, Expectations, and State of the Art. Processes, 9(3): 447, Mar. 2021. ISSN 2227-9717. doi: 10.3390/pr9030447.

44. K.-A. Norton, K. Jin, and A. S. Popel. Modeling triple-negative breast cancer heterogeneity: Effects of stromal macrophages, fibroblasts and tumor vasculature. Journal of Theoretical Biology, 452:56–68, 2018. ISSN 0022-5193. doi: 10.1016/j.jtbi.2018.05.003. URL https://www.sciencedirect.com/science/article/pii/S0022519318302297.

45. K.-A. Norton, C. Gong, S. Jamalian, and A. S. Popel. Multiscale agent-based and hybrid modeling of the tumor immune microenvironment. Processes, 7(1), 2019. ISSN 2227-9717. doi: 10.3390/pr7010037. URL https://www.mdpi.com/2227-9717/7/1/37.

46. L. Ortmann, D. Shi, E. Dassau, F. J. Doyle, B. J. Misgeld, and S. Leonhardt. Automated insulin delivery for type 1 diabetes mellitus patients using gaussian process-based model predictive control. In 2019 American Control Conference (ACC), pages 4118–4123, 2019. doi: 10.23919/ACC.2019.8815258.

47. J. M. Osborne, A. G. Fletcher, J. M. Pitt-Francis, P. K. Maini, and D. J. Gavaghan. Comparing individual-based approaches to modelling the self-organization of multicellular tissues. PLOS Computational Biology, 13(2):e1005387, Feb. 2017. ISSN 1553-7358. doi: 10.1371/journal.pcbi.1005387.

48. G. A. Pavlopoulos, M. Secrier, C. N. Moschopoulos, T. G. Soldatos, S. Kossida, J. Aerts, R. Schneider, and P. G. Bagos. Using graph theory to analyze biological networks. BioData Mining, 4(1):10, Apr. 2011. ISSN 1756-0381. doi: 10.1186/1756-0381-4-10.

49. F. Pedregosa, G. Varoquaux, A. Gramfort, V. Michel, B. Thirion, O. Grisel, M. Blondel, P. Prettenhofer, R. Weiss, V. Dubourg, J. Vanderplas, A. Passos, D. Cournapeau, M. Brucher, M. Perrot, and É. Duchesnay. Scikit-learn: Machine Learning in Python. Journal of Machine Learning Research, 12(85):2825–2830, 2011. ISSN 1533-7928.

50. Q. Peng and F. Vermolen. Agent-based modelling and parameter sensitivity analysis with a finite-element method for skin contraction. Biomech Model Mechanobiol, 19(6):2525–2551, Dec. 2020. ISSN 1617-7940. doi: 10.1007/s10237-020-01354-z.

51. E. E. Peterson, J. M. Ver Hoef, D. J. Isaak, J. A. Falke, M.-J. Fortin, C. E. Jordan, K. McNyset, P. Monestiez, A. S. Ruesch, A. Sengupta, N. Som, E. A. Steel, D. M. Theobald, C. E. Torgersen, and S. J. Wenger. Modelling dendritic ecological networks in space: An integrated network perspective. Ecology Letters, 16(5):707–719, 2013. ISSN 1461-0248. doi: 10.1111/ele.12084.

52. J. Pleyer and C. Fleck. Agent-based models in cellular systems. Frontiers in Physics, 10, 2023. ISSN 2296-424X.

53. A. N. Prybutok, J. Y. Cain, J. N. Leonard, and N. Bagheri. Fighting fire with fire: Deploying complexity in computational modeling to effectively characterize complex biological systems. Current Opinion in Biotechnology, 75:102704, June 2022a. ISSN 0958-1669. doi: 10.1016/j.copbio.2022.102704.

54. A. N. Prybutok, J. S. Yu, J. N. Leonard, and N. Bagheri. Mapping CAR T-Cell Design Space Using Agent-Based Models. Frontiers in Molecular Biosciences, 9, 2022b. ISSN 2296-889X.

55. Z. Z. Shi, C.-H. Wu, and D. Ben-Arieh. Agent-Based Model: A Surging Tool to Simulate Infectious Diseases in the Immune System. Open Journal of Modelling and Simulation, 2014, Jan. 2014. ISSN 2327-4026. doi: 10.4236/ojmsi.2014.21004.

56. E. Sklar. NetLogo, a multi-agent simulation environment. Artif Life, 13(3):303–311, 2007. ISSN 1064-5462. doi: 10.1162/artl.2007.13.3.303.

57. I. M. Sobol. Uniformly distributed sequences with an additional uniform property. USSR Computational Mathematics and Mathematical Physics, 16(5):236–242, Jan. 1976. ISSN 0041-5553. doi: 10.1016/0041-5553(76)90154-3.

58. M. Soheilypour and M. R. K. Mofrad. Agent-Based Modeling in Molecular Systems Biology. BioEssays, 40(7):1800020, 2018. ISSN 1521-1878. doi: 10.1002/bies.201800020.

59. O. Sporns. Structure and function of complex brain networks. Dialogues Clin Neurosci, 15(3):247–262, Sept. 2013. ISSN 1294-8322.

60. M. G. Vander Heiden, L. C. Cantley, and C. B. Thompson. Understanding the Warburg effect: The metabolic requirements of cell proliferation. Science, 324(5930):1029–1033, May 2009. ISSN 1095-9203. doi: 10.1126/science.1160809.

61. I. Vernon, J. Liu, M. Goldstein, J. Rowe, J. Topping, and K. Lindsey. Bayesian uncertainty analysis for complex systems biology models: Emulation, global parameter searches and evaluation of gene functions. BMC Systems Biology, 12(1):1, Jan. 2018. ISSN 1752-0509. doi: 10.1186/s12918-017-0484-3.

62. K. M. Virgilio, K. S. Martin, S. M. Peirce, and S. S. Blemker. Agent-based model illustrates the role of the microenvironment in regeneration in healthy and mdx skeletal muscle. Journal of Applied Physiology, 125(5):1424–1439, 2018.

63. P. Virtanen, R. Gommers, T. E. Oliphant, M. Haberland, T. Reddy, D. Cournapeau, E. Burovski, P. Peterson, W. Weckesser, J. Bright, S. J. van der Walt, M. Brett, J. Wilson, K. J. Millman, N. Mayorov, A. R. J. Nelson, E. Jones, R. Kern, E. Larson, C. J. Carey, I. Polat, Y. Feng, E. W. Moore, J. VanderPlas, D. Laxalde, J. Perktold, R. Cimrman, I. Henriksen, E. A. Quintero, C. R. Harris, A. M. Archibald, A. H. Ribeiro, F. Pedregosa, and P. van Mulbregt. SciPy 1.0: Fundamental algorithms for scientific computing in Python. Nat Methods, 17(3):261–272, Mar. 2020. ISSN 1548-7105. doi: 10.1038/s41592-019-0686-2.

64. Y. Vodovotz and G. An. Agent-based models of inflammation in translational systems biology: A decade later. Wiley Interdisciplinary Reviews: Systems Biology and Medicine, 11(6):e1460, 2019.

65. S. Wang, K. Fan, N. Luo, Y. Cao, F. Wu, C. Zhang, K. A. Heller, and L. You. Massive computational acceleration by using neural networks to emulate mechanism-based biological models. Nat Commun, 10(1):4354, Sept. 2019. ISSN 2041-1723. doi: 10.1038/s41467-019-12342-y.

66. J. West, M. Robertson-Tessi, and A. R. A. Anderson. Agent-based methods facilitate integrative science in cancer. Trends in Cell Biology, 33(4):300–311, Apr. 2023. ISSN 0962-8924, 1879-3088. doi: 10.1016/j.tcb.2022.10.006.

67. J. S. Yu. Arcade. https://github.com/bagherilab/ARCADE, 2023.

68. J. S. Yu and N. Bagheri. Multi-class and multi-scale models of complex biological phenomena. Current Opinion in Biotechnology, 39: 167–173, June 2016. ISSN 0958-1669. doi: 10.1016/j.copbio.2016.04.002.

69. J. S. Yu and N. Bagheri. Agent-Based Models Predict Emergent Behavior of Heterogeneous Cell Populations in Dynamic Microenvironments. Front Bioeng Biotechnol, 8:249, 2020. ISSN 2296-4185. doi: 10.3389/fbioe.2020.00249.

70. J. S. Yu and N. Bagheri. Modular microenvironment components reproduce vascular dynamics de novo in a multi-scale agent-based model. cels, 12(8):795–809.e9, Aug. 2021. ISSN 2405-4712. doi: 10.1016/j.cels.2021.05.007.

