## Supplemental Information for "Incorporating temporal information during feature engineering bolsters emulation of spatio-temporal emergence"

#### Emergent behavior metrics

Emergent behavior metrics are defined and calculated as presented in a previous study of heterogeneous vasculatures in ARCADE (Yu and Bagheri, 2021).

##### Growth rate

Growth rate quantifies the change in colony diameter over time. First, colony diameter is calculated at each time index. Growth rate is the slope of the simple linear regression between $[0, 0.5, \ldots, t_i]$ and corresponding diameters $[D_0, D_{0.5}, \ldots, D_i]$, where $i$ indicates the timepoint. These calculations were performed using the Python function `polyfit` from package `numpy` with degree of 1.

##### Symmetry

Symmetry ($S$) quantifies colony shape at a given timepoint, ranging from 0 (not symmetric) to 1 (perfectly symmetric). For hexagonal coordinates, a colony is perfectly symmetric if for each location (u,v,w), the corresponding five locations (-w,-u,-v), (v,w,u), (-u,-v,-w), (w,u,v), and (-v,-w,-u) are all occupied. Symmetry is calculated as:

$$S = 1 - \frac{1}{N} \sum_{i}^{N} \frac{n_i}{5}$$

where $N$ is the number of unique locations and $n_i$ is the number of corresponding unoccupied locations for a unique location $i$.

##### Activity

Activity ($A$) quantifies the ratio of active (proliferative and migratory) to inactive (necrotic and apoptotic) cells at a given timepoint. This metric ranges between -1 (all non-quiescent cells are inactive) and +1 (all non-quiescent cells are active). An activity value of 0 indicates an equal number of active and inactive cells. Activity is calculated as:

$$A = 2 * \frac{N_a}{N_a + N_i} - 1$$

where $N_a$ is the number of active cells and $N_i$ is the number of inactive cells.

### References

J. S. Yu and N. Bagheri. Modular microenvironment components reproduce vascular dynamics de novo in a multi-scale agent-based model. *cels*, 12(8):795–809.e9, Aug. 2021. ISSN 2405-4712. doi: 10.1016/j.cels.2021.05.007.

**Supp. Table 1.** Topological feature list

| Feature | Code | Equation | Description |
|---|---|---|---|
| Radius | RADIUS | $\frac{\sum_{e \in E} \text{radius of e}}{\|E\|}$ | Average vessel radius |
| Length | LENGTH | $\frac{\sum_{e \in E} \text{length of e}}{\|E\|}$ | Average vessel length |
| Wall | WALL | $\frac{\sum_{e \in E} \text{wall thickness of e}}{\|E\|}$ | Average vessel wall thickness |
| Shear | SHEAR | $\frac{\sum_{e \in E} \text{shear force on e}}{\|E\|}$ | Average vessel shear force |
| Circumference | CIRCUM | $\frac{\sum_{e \in E} \text{circumference of e}}{\|E\|}$ | Average vessel circumference |
| Flow | FLOW | $\frac{\sum_{e \in E} \text{flow through e}}{\|E\|}$ | Average vessel flow |
| Nodes | NODES | $\|V\|$ | Number of nodes in the graph |
| Edges | EDGES | $\|E\|$ | Number of edges in the graph |
| Average eccentricity | AVG_ECCENTRICITY | $\frac{\sum_{v \in V} max(dist(v))}{\|V\|}$ where $dist(v)$ is the shortest distances between $v$ and all other nodes | The average longest shortest path that each nodes is from another node |
| Graph radius | GRADIUS | $min(eccentricity)$ | The minimum eccentricity value |
| Graph diameter | GDIAMETER | $max(eccentricity)$ | The maximum eccentricity value |
| Average shortest path | AVG_SHORTEST_PATH | $\frac{\sum_{v \in V} \frac{(dist(v)}{\|V\|}}{\|V\|} * \frac{1}{\|V\|*(\|V\|-1)}$ where $dist(v)$ is the shortest distances between $v$ and all other nodes | The average shortest path between each nodes and every other node |
| Average in degree | AVG_IN_DEGREES | $\frac{\sum_{v \in V} indegree(v)}{\|V\|}$ | The average in degree in a directed graph |
| Average out degree | AVG_OUT_DEGREES | $\frac{\sum_{v \in V} outdegree(v)}{\|V\|}$ | The average out degree in a directed graph |
| Average degree | AVG_DEGREE | $\frac{\sum_{v \in V} degree(v)}{\|V\|}$ | The average degree in an undirected graph |
| Average clustering | AVG_CLUSTERING | $\frac{\sum_{v \in V} Cl(v)}{\|V\|}$ where $Cl(v)$ is the ratio of the ratio between the number of existing connections between neighbors of $v$ and the maximum possible connections between them | The average clustering coefficient of a graph; the tendency for nodes to form tightly interconnected clusters |
| Average closeness | AVG_CLOSENESS | $\frac{(\|V\|-1)*(\|V\|-1)}{\frac{\sum_{v \in V} dist(v)}{\|V\|}}$ where $dist(v)$ is the shortest distances between $v$ and all other nodes | The average closeness between nodes using the Wasserman and Faust imrpoved formula |
| Average betweenness | AVG_BETWEENNESS | $\frac{\sum_{v \in V} B(v)}{\|V\|} * \frac{1}{(\|V\|-1)*(\|V\|-2)}$ where $B(v)$ is the sum of the fraction of shortest paths passing through $v$ over all pairs of nodes in the graph | The average betweenness of a graph; the average importance of nodes connectors between other nodes in the network |
| Average coreness | AVG_CORENESS | $\frac{\sum_{v \in V} C(v)}{\|V\|}$ where $C(v)$ is the coreness of node $v$ | The average coreness of a graph; the level of connectedness of nodes within a graph |

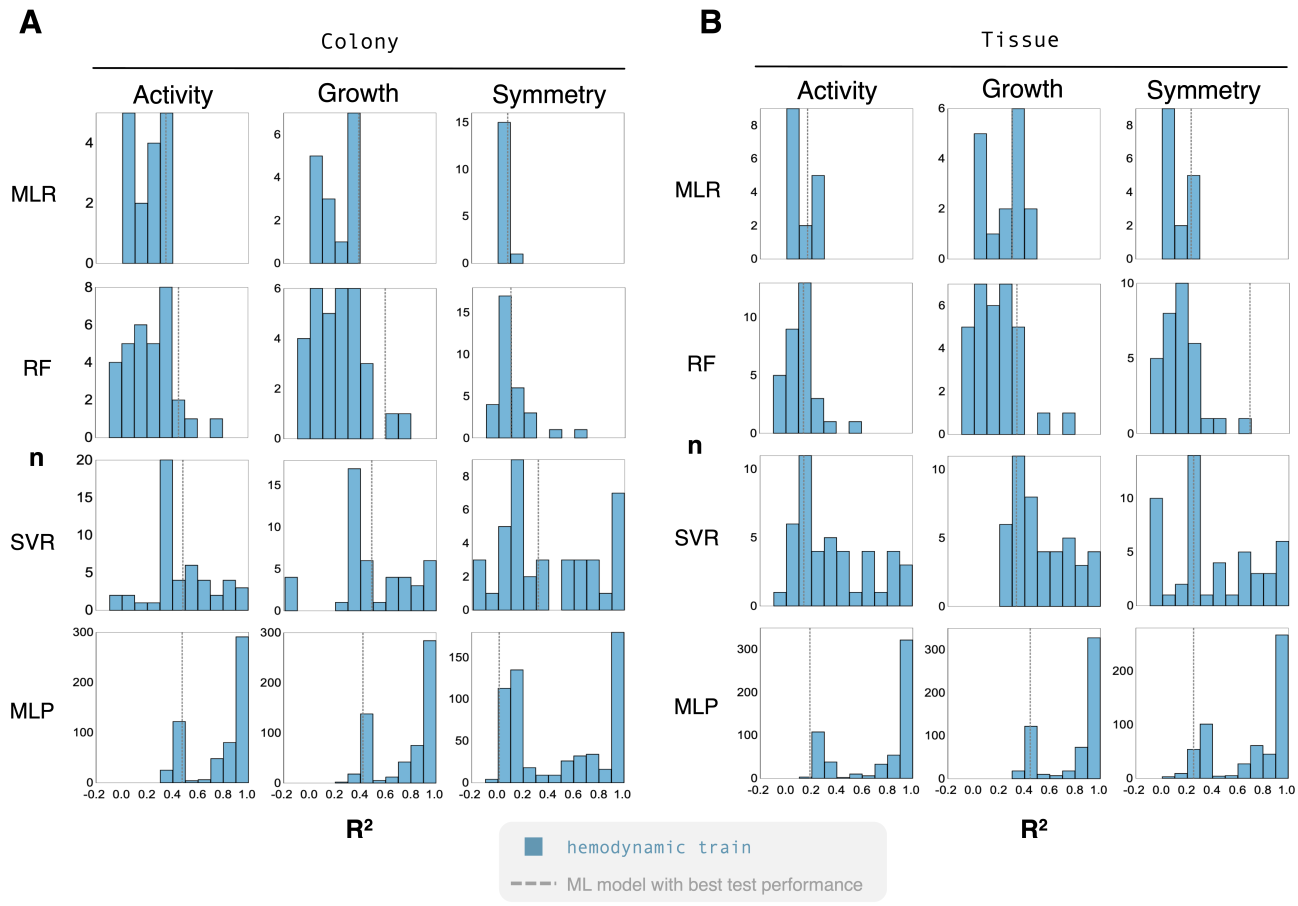


Supp. Fig. 1: **Emulators capture variance across the data during training** — (**A**) Histograms show a range of performances achieved during training for different parameter combinations in the `colony` context. More complex models, such as MLP, can perfectly fit the data in many cases. The vertical dashed line indicates the training performance on model with the best validation performance. (**B**) Similarly, histograms show prediction performance during training across responses and models in the `tissue` context.

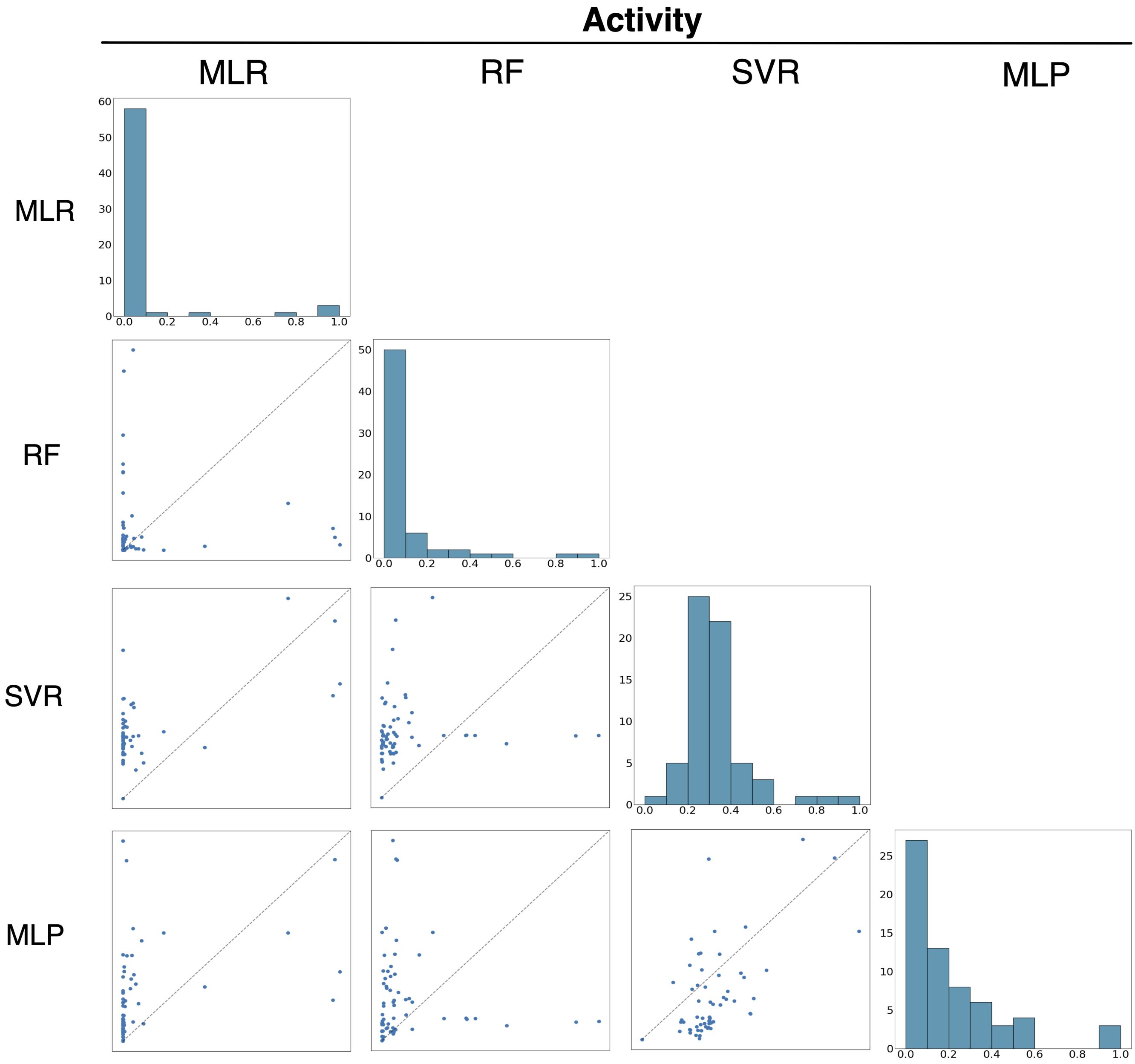


Supp. Fig. 2: **Discrepancy in feature importance across models** — Histograms along the diagonal show the normalized importance distribution of features for each model. Parity plots show the importance of a feature for one model against another. Perfect parity would indicate that the models agreed on the importance level of each feature.

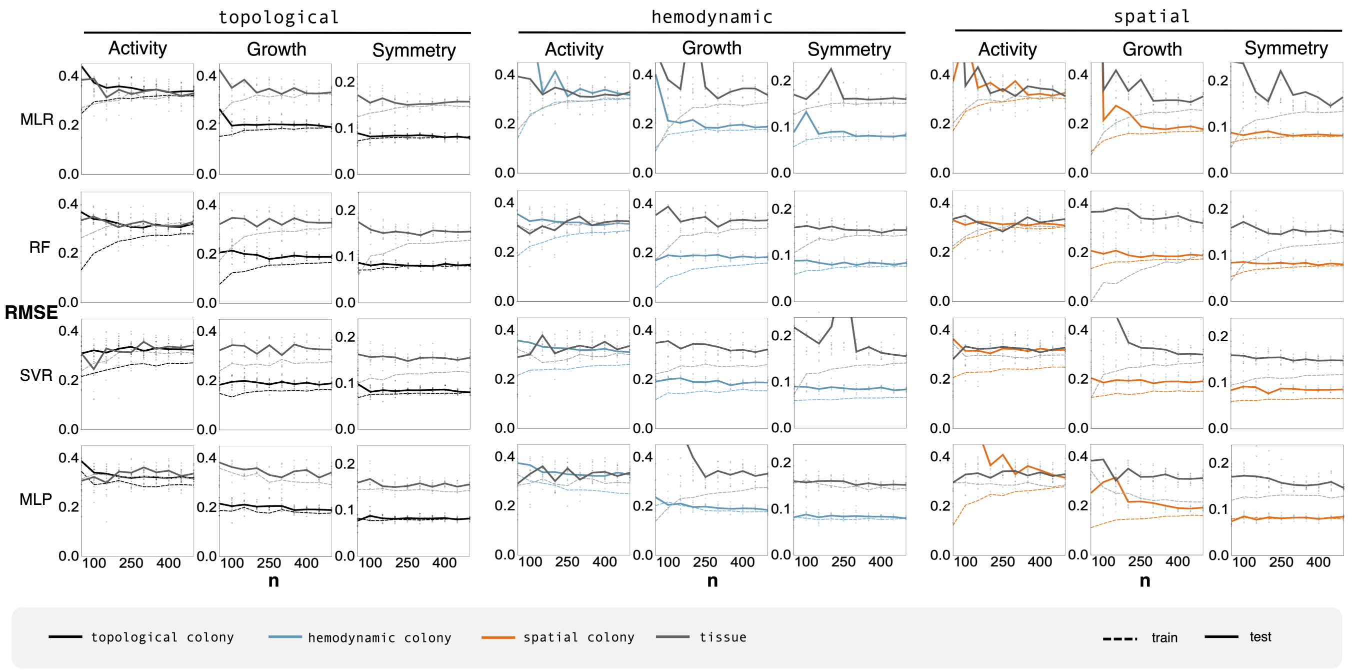


Supp. Fig. 3: **Training data points to diminishing returns** — Line plots indicate the predictive performance of models trained on increasingly large training data sets. In most cases, the RMSE shows diminishing returns of model performance across all feature sets (topolgical, hemodynamic, and spatial), emergent targets (activity, growth, and symmetry), and algorithm (MRL, RF, SVR, MLP).

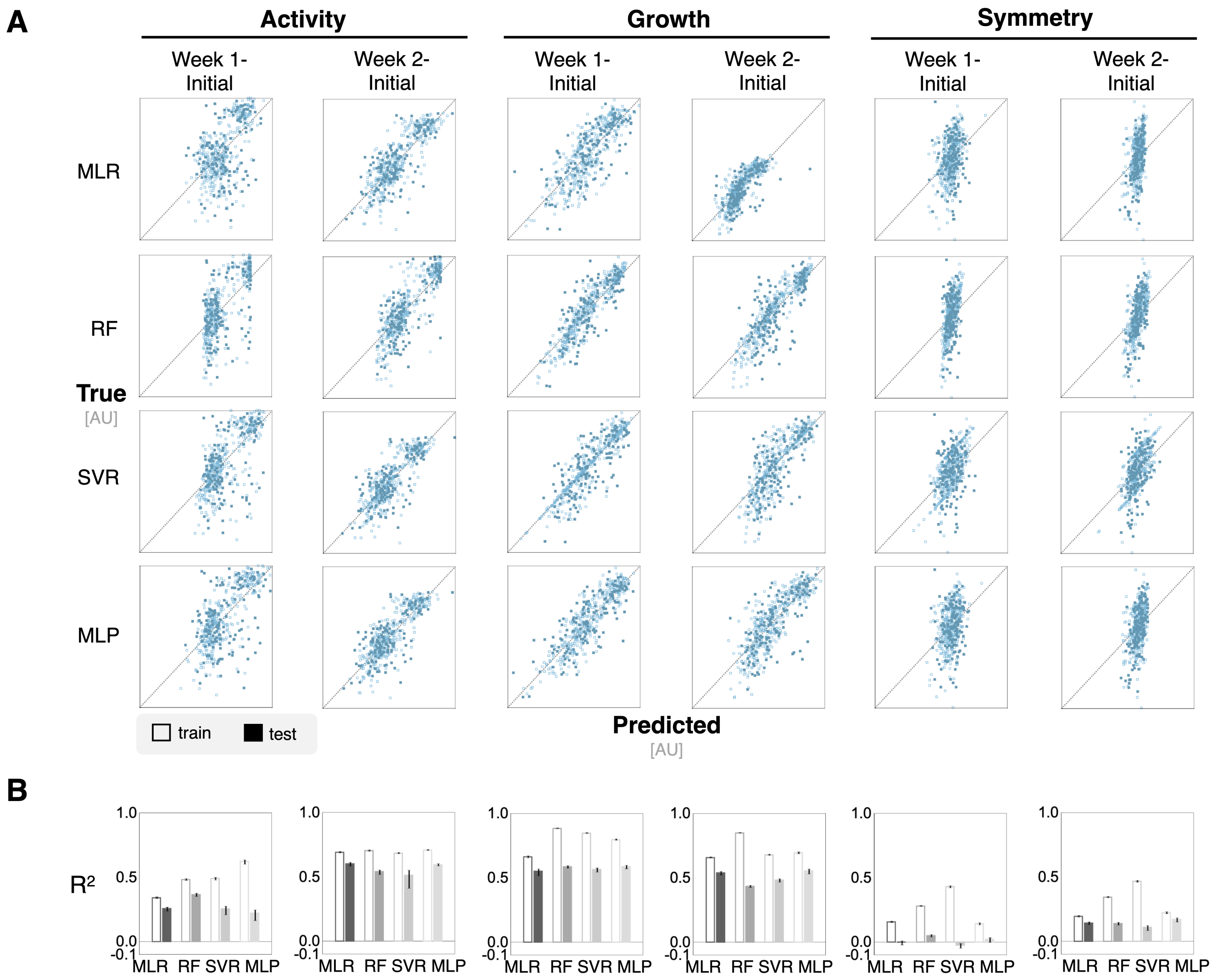


Supp. Fig. 4: **Differential timepoint analysis shows minimal improvement in performance in a `colony` context** — Differential features were calculated by subtracting the features at either one or two weeks from initial features in order to capture the evolution of the features over time. (**A**) Parity plots show the predicted values of emergent targets against the real value to demonstrate fine-grained predictive performance. (**B**) Bar plots show $R^2$ values for ML models trained on differential features. Growth predictions in the `colony` context improve with simulations using week one and initial timepoint differential features.

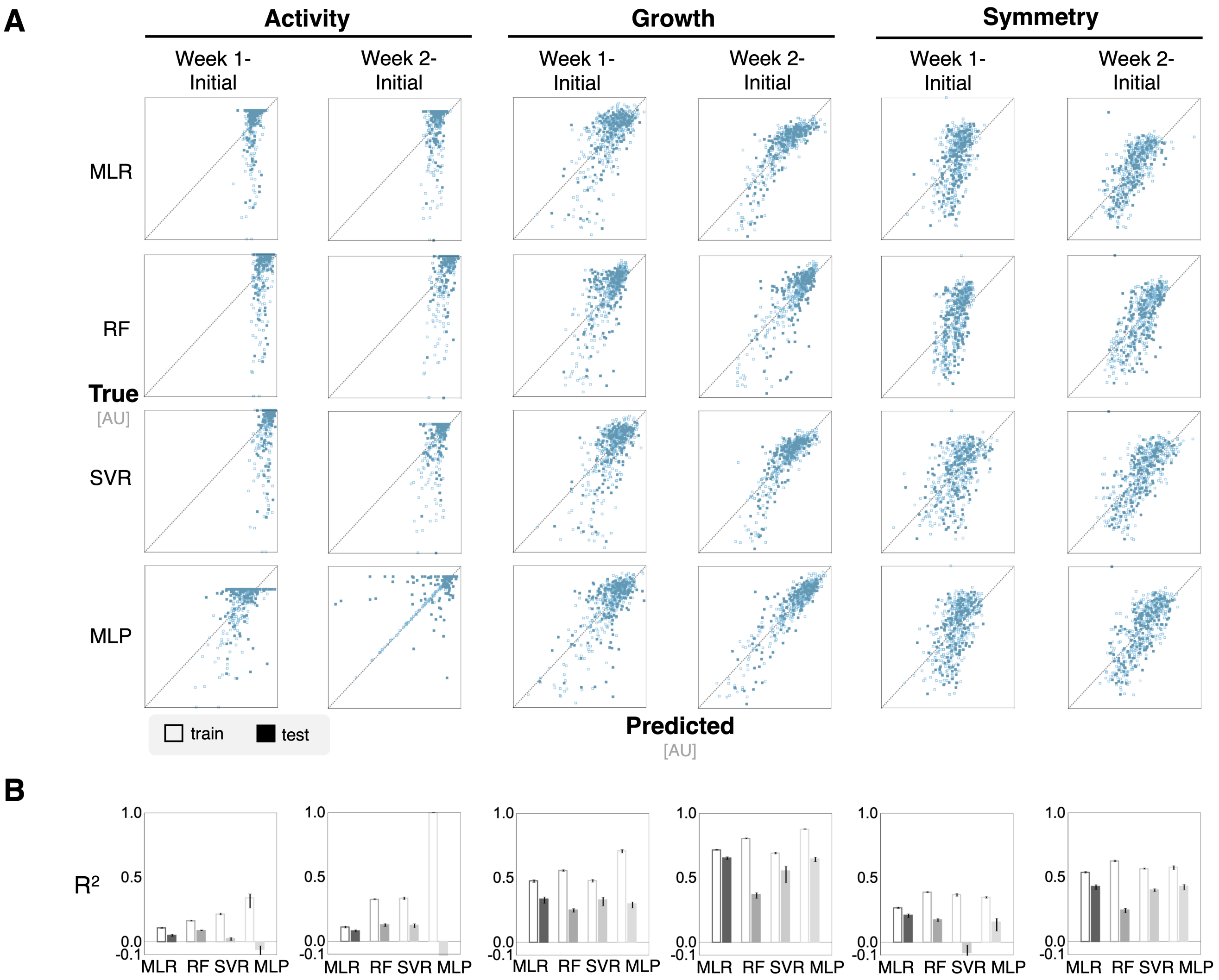


Supp. Fig. 5: **Differential timepoint analysis does not improve performance in a `tissue` context** — Differential features were calculated by subtracting the features at either one or two weeks from initial features in order to capture the evolution of the features over time. (**A**) Parity plots show the predicted values of emergent targets against the real value to demonstrate fine-grained predictive performance. (**B**) Bar plots show $R^2$ values for ML models trained on differential features.

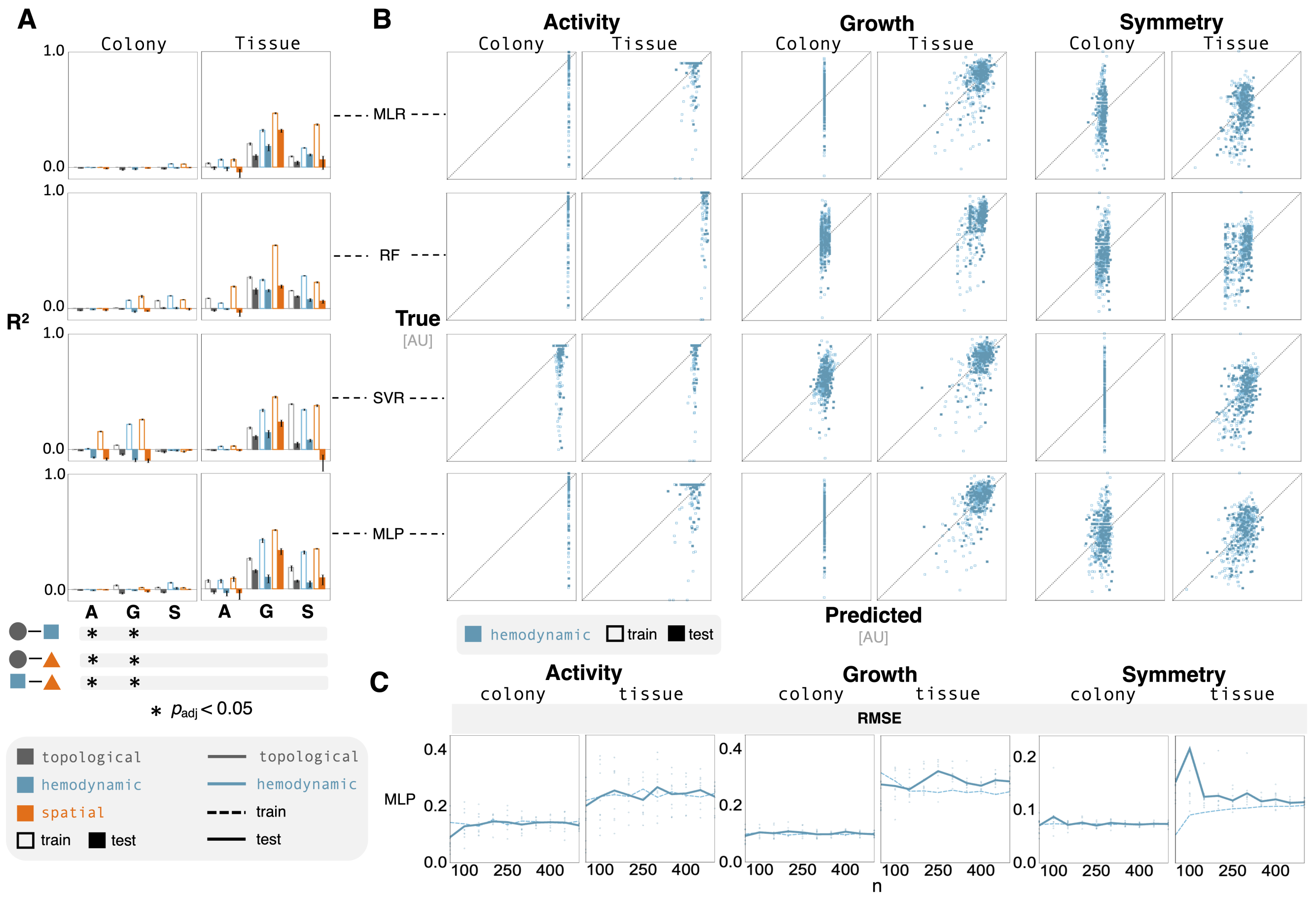


Supp. Fig. 6: **Shortened prediction horizon has minimal effect on emulator performance** — Models predicting spatio-temporal dynamics from after one simulation week. (**A**) Bar plots indicate very poor predictive performance in all cases. Bar chart values range from -0.1 to 1.0; the horizontal axis is at 0.0. The Bonferroni corrected p-values from a two-way ANOVA highlight significant results (noted with black circles) that have an adjusted p-value less that 0.05. (**B**) Parity plots reveal substantial discrepancies in the variance between the predicted and true responses. (**C**) Line plots show predictive performance of the MLP models (the average RMSE) as a function of the size of training data. Performance improvements are limited from additional data. The data points highlight the RMSE from randomized test sets.

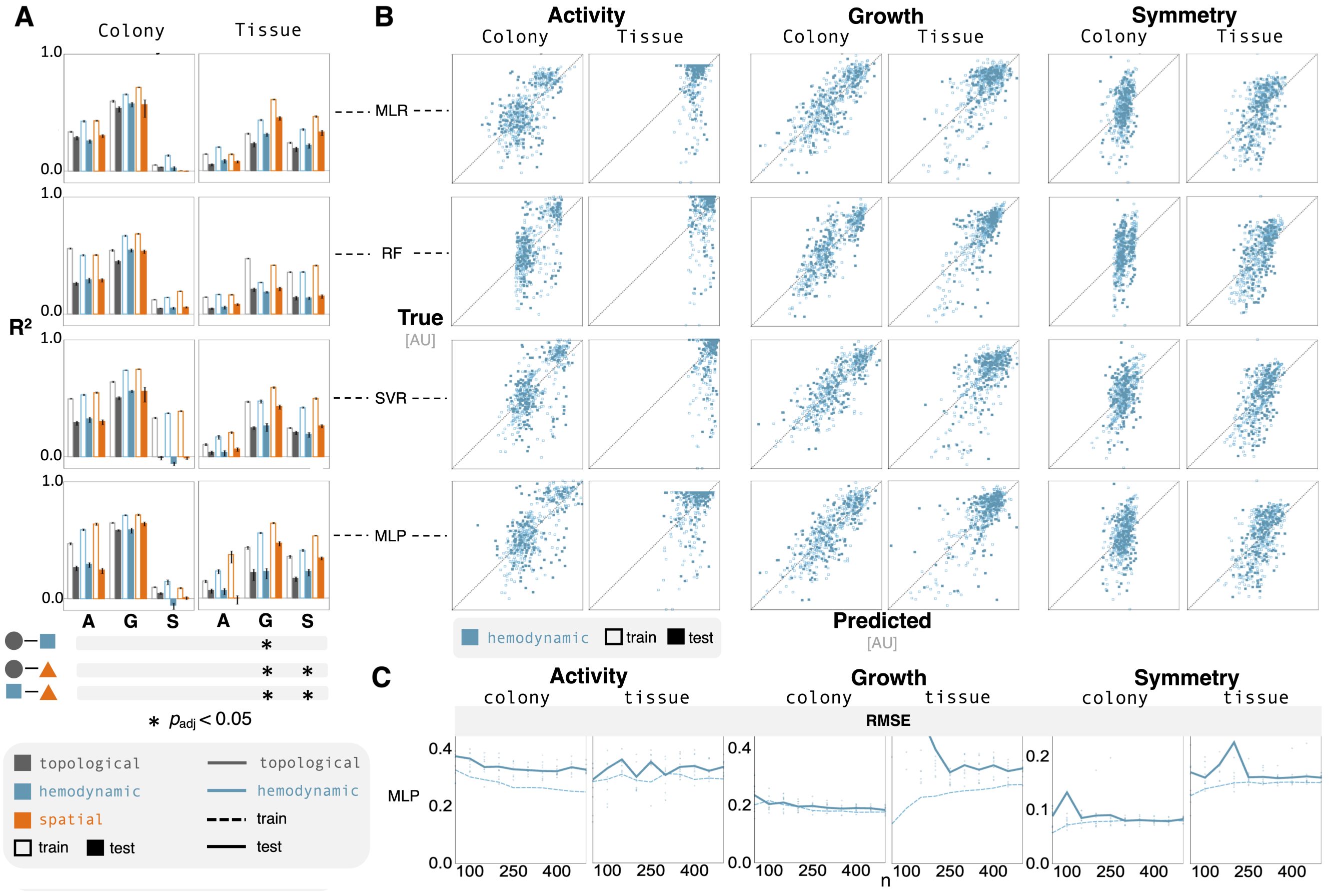


Supp. Fig. 7: **Mid-simulation features show improvement in performance in both contexts** — Models trained on vascular features characterizing network structure at one simulation week result. (**A**) Bar plots show limited improvement from training on mid-simulation features. Bar chart values range from -0.1 to 1.0; the horizontal axis is at 0.0. The Bonferroni corrected p-values from a two-way ANOVA highlight significant results (noted with black circles) that have an adjusted p-value less that 0.05. (**B**) Parity plots reveal large amounts of variance in predicted values with some improvement in growth predictions in the `colony` context. (**C**) Line plots show predictive performance of the MLP models (the average RMSE) as a function of the size of training data. Performance improvements are limited from additional data. The data points highlight the RMSE from randomized test sets.

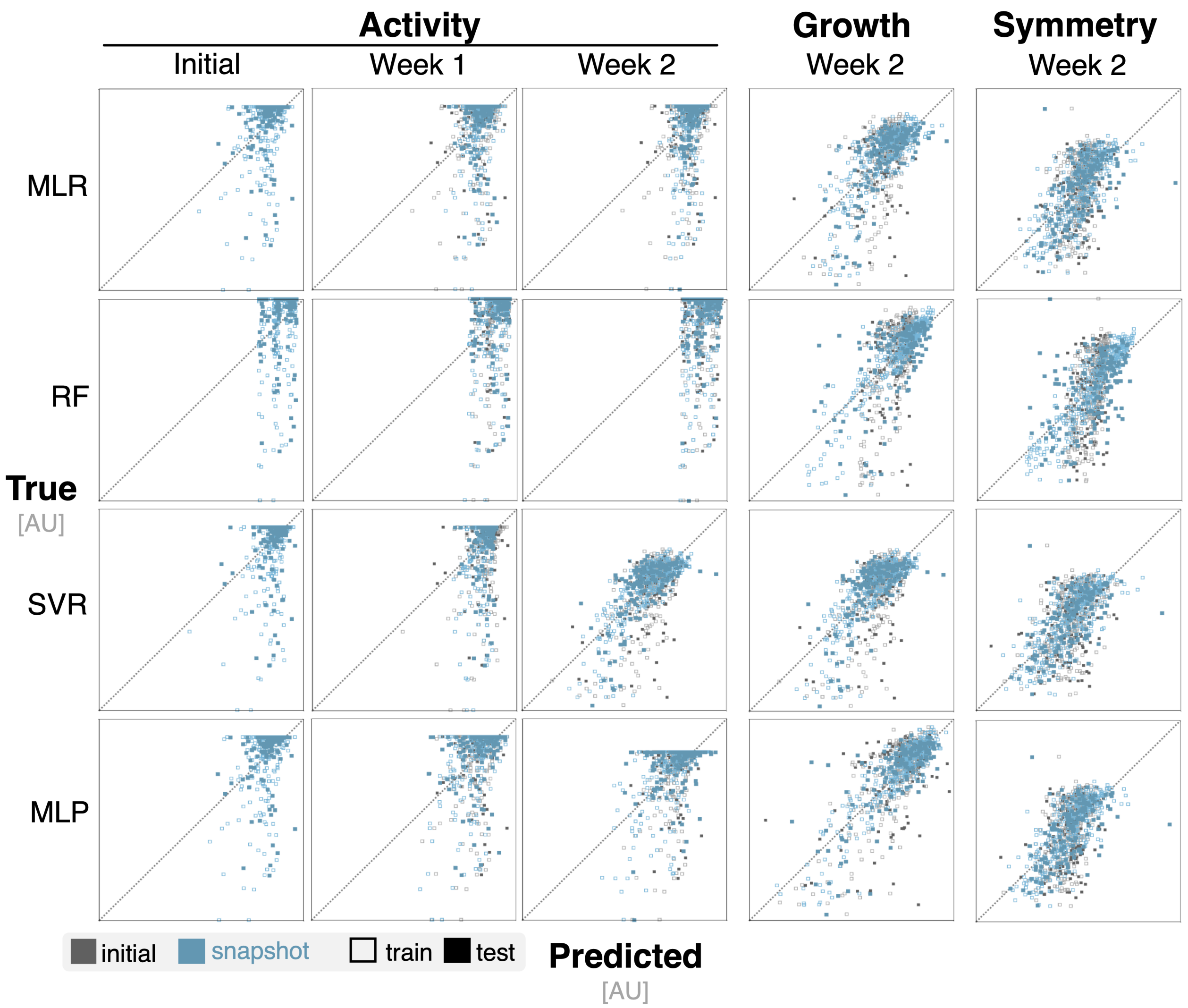


Supp. Fig. 8: **Temporal information improves ML model predictions in `tissue` context** — Parity plots show the predictive performance of ML models trained on features from later timepoints against emulators trained on features from the initial timepoint. Improved prediction of activity is limited; growth and symmetry reflect minimal improvements.

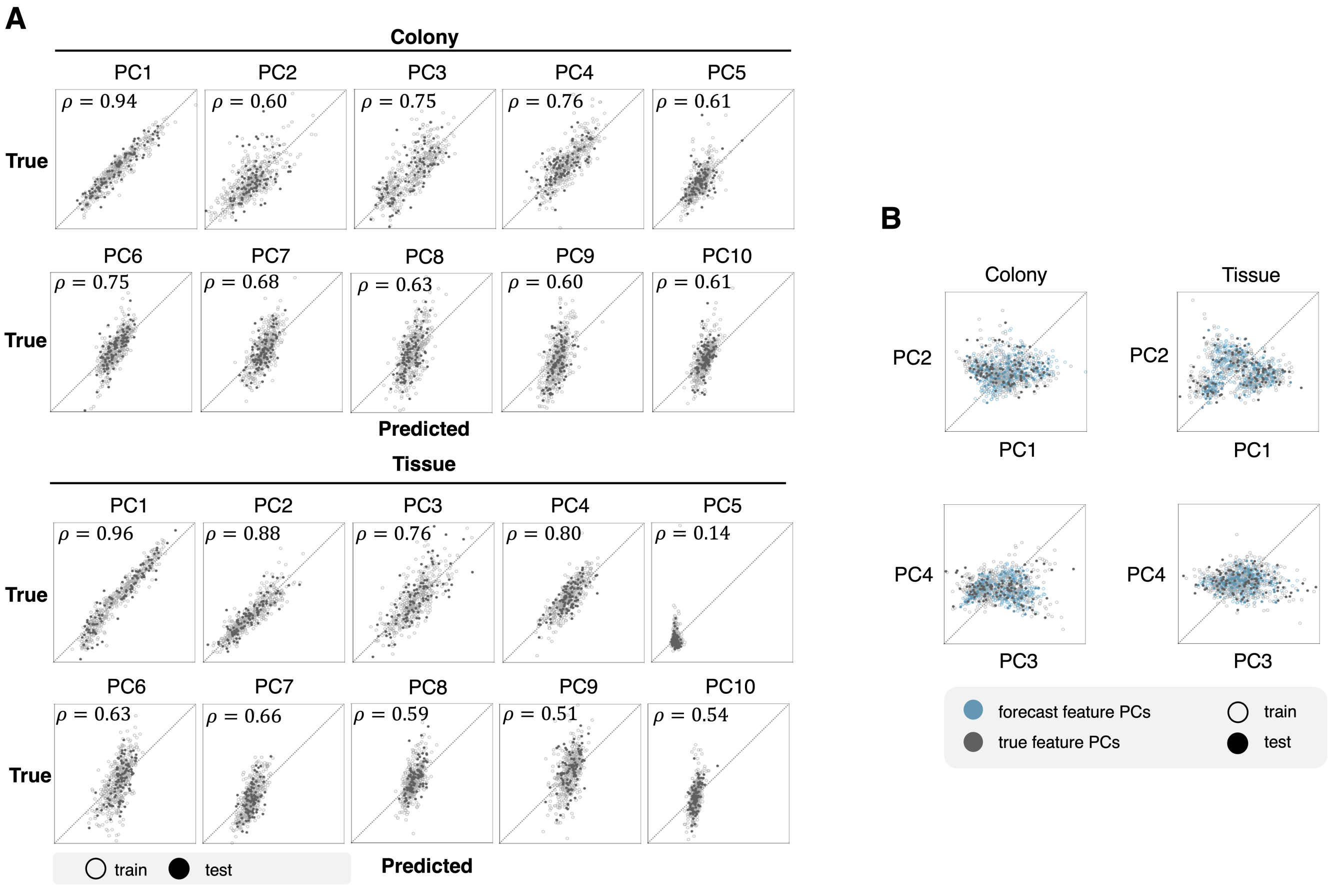


Supp. Fig. 9: **RNN model captures vascular feature variance** — (**A**) Parity plots highlight the performance of a RNN model at predicting network metric features. Both the simulated features and forecasted features were combined to perform PCA. The dimensionality of the feature set are reduced to the first 10 principal components, which represent 95% of the feature variance. The Pearson correlation coefficient ($\rho$) is reported for each parity plot. (**B**) Scatter plots illustrate the overlap between true and forecasted features using the first four principal components.

**Supp. Table 2.** Hemodynamic feature list

| Feature | Code |
|---|---|
| Graph radius weighted by flow through the vessel | GRADIUS:FLOW |
| Graph diameter weighted by flow | GDIAMETER:FLOW |
| Average eccentricity weighted by flow | AVG_ECCENTRICITY:FLOW |
| Average shortest path weighted by flow | AVG_SHORTEST_PATH:FLOW |
| Average closeness weighted by flow | AVG_CLOSENESS:FLOW |
| Average betweenness weighted by flow | AVG_BETWEENNESS:FLOW |
| Graph radius weighted by vessel wall thickness | GRADIUS:WALL |
| Graph diameter weighted by wall | GDIAMETER:WALL |
| Average eccentricity weighted by wall | AVG_ECCENTRICITY:WALL |
| Average shortest path weighted by wall | AVG_SHORTEST_PATH:WALL |
| Average closeness weighted by wall | AVG_CLOSENESS:WALL |
| Average betweenness weighted by wall | AVG_BETWEENNESS:WALL |
| Graph radius weighted by vessel shear force | GRADIUS:SHEAR |
| Graph diameter weighted by shear | GDIAMETER:SHEAR |
| Average eccentricity weighted by shear | AVG_ECCENTRICITY:SHEAR |
| Average shortest path weighted by shear | AVG_SHORTEST_PATH:SHEAR |
| Average closeness weighted by shear | AVG_CLOSENESS:SHEAR |
| Average betweenness weighted by shear | AVG_BETWEENNESS:SHEAR |
| Graph radius weighted by vessel radius | GRADIUS:RADIUS |
| Graph diameter weighted by radius | GDIAMETER:RADIUS |
| Average eccentricity weighted by radius | AVG_ECCENTRICITY:RADIUS |
| Average shortest path weighted by radius | AVG_SHORTEST_PATH:RADIUS |
| Average closeness weighted by radius | AVG_CLOSENESS:RADIUS |
| Average betweenness weighted by radius | AVG_BETWEENNESS:RADIUS |
| Graph radius weighted by average pressure across each vessel | GRADIUS:PRESSURE_AVG |
| Graph diameter weighted by average pressure | GDIAMETER:PRESSURE_AVG |
| Average eccentricity weighted by average pressure | AVG_ECCENTRICITY:PRESSURE_AVG |
| Average shortest path weighted by average pressure | AVG_SHORTEST_PATH:PRESSURE_AVG |
| Average closeness weighted by average pressure | AVG_CLOSENESS:PRESSURE_AVG |
| Average betweenness weighted by average pressure | AVG_BETWEENNESS:PRESSURE_AVG |
| Graph radius weighted by pressure delta across each vessel | GRADIUS:PRESSURE_DELTA |
| Graph diameter weighted by pressure delta | GDIAMETER:PRESSURE_DELTA |
| Average eccentricity weighted by pressure delta | AVG_ECCENTRICITY:PRESSURE_DELTA |
| Average shortest path weighted by pressure delta | AVG_SHORTEST_PATH:PRESSURE_DELTA |
| Average closeness weighted by pressure delta | AVG_CLOSENESS:PRESSURE_DELTA |
| Average betweenness weighted by pressure delta | AVG_BETWEENNESS:PRESSURE_DELTA |
| Graph radius weighted by average oxygen across each vessel | GRADIUS:OXYGEN_AVG |
| Graph diameter weighted by average oxygen | GDIAMETER:OXYGEN_AVG |
| Average eccentricity weighted by average oxygen | AVG_ECCENTRICITY:OXYGEN_AVG |
| Average shortest path weighted by average oxygen | AVG_SHORTEST_PATH:OXYGEN_AVG |
| Average closeness weighted by average oxygen | AVG_CLOSENESS:OXYGEN_AVG |
| Average betweenness weighted by average oxygen | AVG_BETWEENNESS:OXYGEN_AVG |
| Graph radius weighted by oxygen delta across each vessel | GRADIUS:OXYGEN_DELTA |
| Graph diameter weighted by oxygen delta | GDIAMETER:OXYGEN_DELTA |
| Average eccentricity weighted by oxygen delta | AVG_ECCENTRICITY:OXYGEN_DELTA |
| Average shortest path weighted by oxygen delta | AVG_SHORTEST_PATH:OXYGEN_DELTA |
| Average closeness weighted by oxygen delta | AVG_CLOSENESS:OXYGEN_DELTA |
| Average betweenness weighted by oxygen delta | AVG_BETWEENNESS:OXYGEN_DELTA |

**Supp. Table 3.** Spatial feature list

| Feature | Code |
|---|---|
| Graph radius weighted by inverse distance | GRADIUS:INVERSE_DISTANCE |
| Graph diameter weighted by inverse distance | GDIAMETER:INVERSE_DISTANCE |
| Average eccentricity weighted by inverse distance | AVG_ECCENTRICITY:INVERSE_DISTANCE |
| Average shortest path weighted by inverse distance | AVG_SHORTEST_PATH:INVERSE_DISTANCE |
| Average closeness weighted by inverse distance | AVG_CLOSENESS:INVERSE_DISTANCE |
| Average betweenness weighted by inverse distance | AVG_BETWEENNESS:INVERSE_DISTANCE |
| Average eccentricity weighted by distance | AVG_ECCENTRICITY_WEIGHTED |
| Average closeness weighted by distance | AVG_CLOSENESS_WEIGHTED |
| Average coreness weighted by distance | AVG_CORENESS_WEIGHTED |
| Average betweenness weighted by distance | AVG_BETWEENNESS_WEIGHTED |
| Average in degree weighted by distance | AVG_IN_DEGREES_WEIGHTED |
| Average out degree weighted by distance | AVG_OUT_DEGREES_WEIGHTED |
| Average degree weighted by distance | AVG_DEGREE_WEIGHTED |

**Supp. Table 4.** Emulator run times across feature sets on an m5.large EC2 instance

| Model | Feature set | Time (m) ± S.D. |
|---|---|---|
| MLR | Topological | 0.21 ± 0.01 |
| | Hemodynamic | 1.45 ± 0.03 |
| | Spatial | 1.63 ± 0.04 |
| RF | Topological | 1.11 ± 0.01 |
| | Hemodynamic | 2.51 ± 0.26 |
| | Spatial | 2.78 ± 0.08 |
| SVR | Topological | 1.07 ± 0.01 |
| | Hemodynamic | 3.62 ± 0.06 |
| | Spatial | 3.68 ± 0.05 |
| MLP | Topological | 128.62 ± 1.61 |
| | Hemodynamic | 285.71 ± 1.18 |
| | Spatial | 286.20 ± 0.77 |

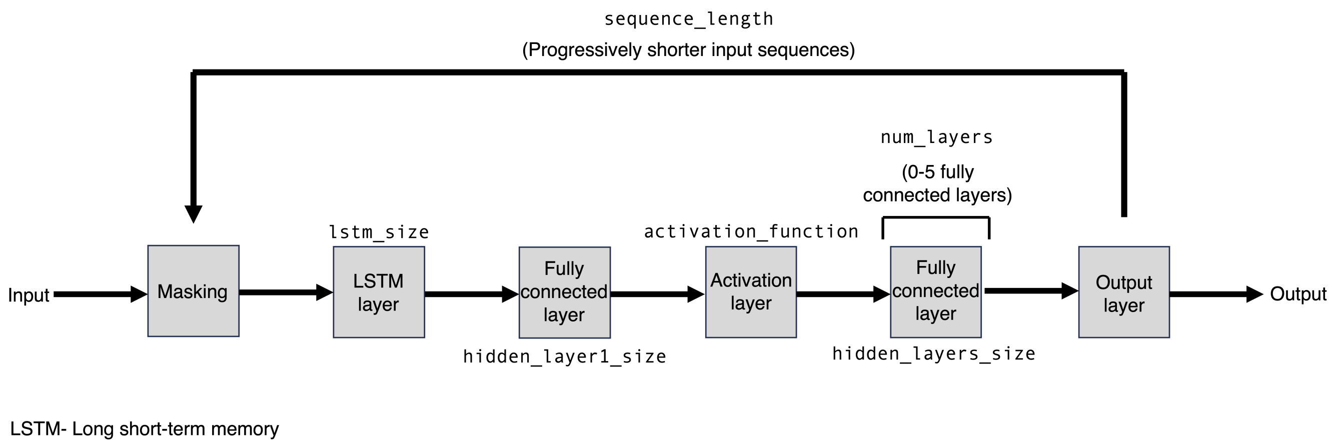


Supp. Fig. 10: **Architecture and structure of the RNN used for feature prediction** — This flowchart describes the layers used while training the feature prediction RNN. All parameter values are defined in Supp. Table 7 and optimal values are provided in Supp. Table 8.

**Supp. Table 5.** Top performing hyperparameters for emulation models in a colony context

| Model | Emergent Target | Feature set | Hyperparameters |
|---|---|---|---|
| MLR | Activity | Topological | alpha:0.001;l1_ratio:0.1;max_iter:10000 |
| | | Hemodynamic | alpha:0.0023; l1_ratio:0.6625; max_iter:10000 |
| | | Spatial | alpha:0.0015; l1_ratio:0.9437; max_iter:10000 |
| | Growth | Topological | alpha:0.001; l1_ratio:0.1 ; max_iter:10000 |
| | | Hemodynamic | alpha:0.0036; l1_ratio:0.3812; max_iter:10000 |
| | | Spatial | alpha:0.0036; l1_ratio:0.3812; max_iter:10000 |
| | Symmetry | Topological | alpha:0.0056; l1_ratio:0.775; max_iter:10000 |
| | | Hemodynamic | alpha:0.0036; l1_ratio:0.3812; max_iter:10000 |
| | | Spatial | alpha:0.0316; l1_ratio:0.55; max_iter:10000 |
| RF | Activity | Topological | n_estimators:13; max_features:0.1778; max_depth:38; min_samples_split:0.0177; min_samples_leaf:0.0177; bootstrap:True |
| | | Hemodynamic | n_estimators:81; max_features:0.2371; max_depth:81; min_samples_split:0.0133; min_samples_leaf:0.0749; bootstrap:False |
| | | Spatial | n_estimators:81; max_features:0.2371; max_depth:81; min_samples_split:0.0133; min_samples_leaf:0.0749; bootstrap:False |
| | Growth | Topological | n_estimators:26; max_features:0.3162; max_depth:75; min_samples_split:0.3162; min_samples_leaf:0.0316; bootstrap:False |
| | | Hemodynamic | n_estimators:81; max_features:0.2371; max_depth:81; min_samples_split:0.0133; min_samples_leaf:0.0749; bootstrap:False |
| | | Spatial | n_estimators:81; max_features:0.2371; max_depth:81; min_samples_split:0.0133; min_samples_leaf:0.0749; bootstrap:False |
| | Symmetry | Topological | n_estimators:57; max_features:0.0749; max_depth:7; min_samples_split:0.4216; min_samples_leaf:0.0237; bootstrap:False |
| | | Hemodynamic | n_estimators:20; max_features:0.0421; max_depth:94; min_samples_split:0.0749; min_samples_leaf:0.1333; bootstrap:False |
| | | Spatial | n_estimators:94; max_features:0.0133; max_depth:69; min_samples_split:0.0237; min_samples_leaf:0.04216; bootstrap:True |
| SVR | Activity | Topological | C:0.5623; epsilon:0.1778; kernel:rbf |
| | | Hemodynamic | C:0.5623; epsilon:0.1778; kernel:rbf |
| | | Spatial | C:0.5623; epsilon:0.1778; kernel:rbf |
| | Growth | Topological | C:0.1; epsilon:0.001; kernel:rbf |
| | | Hemodynamic | C:0.2371; epsilon:0.0749; kernel:rbf |
| | | Spatial | C:0.2371; epsilon:0.0749; kernel:linear |
| | Symmetry | Topological | C:0.2371; epsilon:0.0749; kernel:rbf |
| | | Hemodynamic | C:0.2371; epsilon:0.07498; kernel:rbf |
| | | Spatial | C:0.2371; epsilon:0.0749; kernel:rbf |
| MLP | Activity | Topological | alpha:0.3162; activation:logistic; hidden_layer_sizes:[50, 25]; solver:lbfgs; max_iter:1000 |
| | | Hemodynamic | alpha:0.5623; activation:logistic; hidden_layer_sizes:[25, 50]; solver:lbfgs; max_iter:1000 |
| | | Spatial | alpha:0.5623; activation:logistic; hidden_layer_sizes:[25, 50]; solver:lbfgs; max_iter:1000 |
| | Growth | Topological | alpha:0.1778; activation:logistic; hidden_layer_sizes:[25, 25]; solver:lbfgs; max_iter:1000 |
| | | Hemodynamic | alpha:0.5623; activation:logistic; hidden_layer_sizes:[5]; solver:lbfgs; max_iter:1000 |
| | | Spatial | alpha:0.3162; activation:logistic; hidden_layer_sizes:[25, 25]; solver:lbfgs; max_iter:1000 |
| | Symmetry | Topological | alpha:0.3162; activation:logistic; hidden_layer_sizes:[50]; solver:lbfgs; max_iter:1000 |
| | | Hemodynamic | alpha:0.0005; activation:identity; hidden_layer_sizes:[5, 10]; solver:lbfgs; max_iter:1000 |
| | | Spatial | alpha:0.1778; activation:logistic; hidden_layer_sizes:[50, 50]; solver:lbfgs; max_iter:1000 |

**Supp. Table 6.** Top performing hyperparameters for emulation models in a `tissue` context

| Model | Emergent Target | Feature set | Hyperparameters |
|---|---|---|---|
| MLR | Activity | Topological | alpha:0.0086; l1_ratio:0.2687; max_iter:10000 |
| | | Hemodynamic | alpha:0.0056; l1_ratio:0.775; max_iter:10000 |
| | | Spatial | alpha:0.0749; l1_ratio:0.2125; max_iter:10000 |
| | Growth | Topological | alpha:0.0056; l1_ratio:0.775; max_iter:10000 |
| | | Hemodynamic | alpha:0.0133; l1_ratio:0.4375; max_iter:10000 |
| | | Spatial | alpha:0.0023; l1_ratio:0.6625; max_iter:10000 |
| | Symmetry | Topological | alpha:0.0015; l1_ratio:0.9437; max_iter:10000 |
| | | Hemodynamic | alpha:0.0015; l1_ratio:0.9437; max_iter:10000 |
| | | Spatial | alpha:0.0015; l1_ratio:0.9437; max_iter:10000 |
| RF | Activity | Topological | n_estimators:38; max_features:0.0562; max_depth:63; min_samples_split:0.5623; min_samples_leaf:0.0562; bootstrap:False |
| | | Hemodynamic | n_estimators:38; max_features:0.0562; max_depth:63; min_samples_split:0.5623; min_samples_leaf:0.0562; bootstrap:False |
| | | Spatial | n_estimators:20; max_features:0.0421; max_depth:94; min_samples_split:0.0749; min_samples_leaf:0.1333; bootstrap:False |
| | Growth | Topological | n_estimators:13; max_features:0.1778; max_depth:38; min_samples_split:0.0177; min_samples_leaf:0.0177; bootstrap:True |
| | | Hemodynamic | n_estimators:57; max_features:0.0749; max_depth:7; min_samples_split:0.4216; min_samples_leaf:0.0237; bootstrap:True |
| | | Spatial | n_estimators:81; max_features:0.2371; max_depth:81; min_samples_split:0.0133; min_samples_leaf:0.0749; bootstrap:False |
| | Symmetry | Topological | n_estimators:94; max_features:0.0133; max_depth:69; min_samples_split:0.0237; min_samples_leaf:0.0421; bootstrap:False |
| | | Hemodynamic | n_estimators:94; max_features:0.0133; max_depth:69; min_samples_split:0.0237; min_samples_leaf:0.0421; bootstrap:False |
| | | Spatial | n_estimators:81; max_features:0.2371; max_depth:81; min_samples_split:0.0133; min_samples_leaf:0.0749; bootstrap:False |
| SVR | Activity | Topological | C:0.5623; epsilon:0.1778; kernel:linear |
| | | Hemodynamic | C:0.2371; epsilon:0.0749; kernel:linear |
| | | Spatial | C:0.2371; epsilon:0.0749; kernel:rbf |
| | Growth | Topological | C:0.5623; epsilon:0.1778; kernel:rbf |
| | | Hemodynamic | C:1.333; epsilon:0.0133; kernel:linear |
| | | Spatial | C:4.8696; epsilon:0.0205; kernel:linear |
| | Symmetry | Topological | C:0.2371; epsilon:0.0749; kernel:linear |
| | | Hemodynamic | C:0.2371; epsilon:0.0749; kernel:linear |
| | | Spatial | C:0.1; epsilon:0.001; kernel:linear |
| MLP | Activity | Topological | alpha:0.3162; activation:logistic; hidden_layer_sizes:[50, 25]; solver:lbfgs; max_iter:1000 |
| | | Hemodynamic | alpha:0.5623; activation:logistic; hidden_layer_sizes:[50, 25]; solver:lbfgs; max_iter:1000 |
| | | Spatial | alpha:0.5623; activation:logistic; hidden_layer_sizes:[50, 25]; solver:lbfgs; max_iter:1000 |
| | Growth | Topological | alpha:0.5623; activation:logistic; hidden_layer_sizes:[50]; solver:lbfgs; max_iter:1000 |
| | | Hemodynamic | alpha:0.5623; activation:identity; hidden_layer_sizes:[25]; solver:lbfgs; max_iter:1000 |
| | | Spatial | alpha:0.5623; activation:identity; hidden_layer_sizes:[5]; solver:lbfgs; max_iter:1000 |
| | Symmetry | Topological | alpha:0.5623; activation:tanh; hidden_layer_sizes:[50, 50]; solver:lbfgs; max_iter:1000 |
| | | Hemodynamic | alpha:0.5623; activation:identity; hidden_layer_sizes:[5, 5]; solver:lbfgs; max_iter:1000 |
| | | Spatial | alpha:0.5623; activation:identity; hidden_layer_sizes:[5, 10]; solver:lbfgs; max_iter:1000 |

**Supp. Table 7.** Hyperparameters used in RNN architecture grid search

| Hyperparameter | Code | Potential value |
| --- | --- | --- |
| Training sequence lengths | sequence_length | [15, 10, 5, 3, 1]] |
| | | [15, 10, 5, 1] |
| | | [15, 5, 3, 1] |
| LSTM layer size | lstm_size | 64 |
| | | 128 |
| | | 256 |
| | | 512 |
| | | 1024 |
| Size of first hidden layer | hidden_layer1_size | 64 |
| | | 128 |
| | | 256 |
| | | 512 |
| | | 1024 |
| Size of the remaining hidden layers | hidden_layers_size | 64 |
| | | 128 |
| | | 256 |
| | | 512 |
| | | 1024 |
| Number of hidden layers after the first | num_layers | 0 |
| | | 1 |
| | | 2 |
| | | 3 |
| | | 4 |
| | | 5 |
| Activation function after first hidden layer | activation_function | `relu` |
| | | `tanh` |

**Supp. Table 8.** Top performing architecture parameters for RNN

| Context | Parameter | Value |
| --- | --- | --- |
| colony | `sequence_length` | [15, 10, 5, 3, 1] |
| | `lstm_size` | 512 |
| | `hidden_layer1_size` | 1024 |
| | `hidden_layers_size` | 1024 |
| | `num_layers` | 2 |
| | `activation_function` | `relu` |
| tissue | `sequence_length` | [15, 10, 5, 3, 1] |
| | `lstm_size` | 512 |
| | `hidden_layer1_size` | 512 |
| | `hidden_layers_size` | 1024 |
| | `num_layers` | 1 |
| | `activation_function` | `relu` |
